# Estimates of molecular convergence reveal genes with intermediate pleiotropy underlying adaptive variation across teleost fish

**DOI:** 10.1101/2024.06.24.600426

**Authors:** Agneesh Barua, Malvika Srivastava, Brice Beinsteiner, Vincent Laudet, Marc Robinson-Rechavi

## Abstract

Molecular convergence, where specific non-synonymous changes in protein-coding genes lead to identical amino acid substitutions across multiple lineages, provides strong evidence of adaptive evolution. Detecting this signal across diverse taxa can reveal broad evolutionary mechanisms that may not be apparent when studying individual lineages. In this study, we search for convergent substitutions in the most speciose group of vertebrates, teleost fishes. Using an unsupervised approach, we detected convergence in 89 protein-coding genes across 143 chromosomal-level genomes. To assess their functional implications, we integrate data on protein properties, gene expression across species and tissues, single-cell RNA sequencing of zebrafish embryonic development, and gene perturbation experiments in zebrafish. The convergent genes were associated with diverse processes including embryonic development, tissue morphogenesis, metabolism, and responses to hormones and heat stress. The convergent substitutions altered amino acid properties, with some occurring at functionally critical sites. Notably, only one-third of these genes were tissue-specific, while the majority were expressed across multiple tissues and cell types. Genetic perturbation data further showed that the convergent genes can affect multiple structures across diverse tissues. These results highlight the pleiotropic nature of the convergent genes. Using an evolutionary modeling approach, we show that adaptive variation tends to accumulate in genes with intermediate pleiotropy, enabling organisms to overcome selective pressures during ecological shifts. Although traditionally considered a source of constraint, we argue that adaptation via genes with intermediate pleiotropy might be particularly advantageous following periods of ecological shifts, and can potentially lead to the evolution of convergent phenotypes.

## Introduction

Evolutionary convergence, characterised by different lineages independently acquiring similar character states, in response to similar ecological challenges, is pervasive in nature ^1^. While convergence typically invokes images of recurring phenotypic traits, convergence can also occur at the molecular level where the same amino acid substitution can occur at the same site in orthologous proteins ^2,3^. Identifying molecular signatures of convergence is integral to understanding the genetic basis behind the evolution of phenotypes, as well as aiding in recognising adaptive developmental processes across diverse groups of animals ^4,5^. Indeed, studies of molecular convergence have provided insights into the origin of traits like echolocation in birds and mammals, toxin resistance in beetles, and carnivory in plants, just to name a few ^6–8^. The earliest methods to estimate molecular convergence relied on reconstructed ancestral sequences to identify sites in foreground branches that had the same amino acid substitution but differed in their ancestral sequence ^9,10^. Later methods refined this approach by estimating ratios of repeated and divergent changes in amino acids in foreground branches to distinguish true instances of repeated substitutions from null expectations ^11,12^. However, requiring foreground lineages to share the same amino acid substitution can limit the detection of functionally convergent changes that arise through different amino acid substitutions ^13,14^. Additionally, restricting analyses to a predefined set of foreground branches based on an *a priori* phenotype of interest may overlook other potential adaptive changes. To capture a broader range of repeated evolution, studies would benefit from unbiased analytical approaches.

The prevalence of evolutionary convergence highlights a deterministic aspect of the evolutionary process, leading some researchers to propose that evolution may, to some extent, be predictable ^15^. Studies have consistently identified certain gene categories—such as gas transporters, sensory proteins, and xenobiotic-processing enzymes—as frequent targets of adaptation to new environments ^16^. However, one class of genes has long been considered poor candidates for adaptive evolution: pleiotropic genes. Pleiotropy is the property of genes or mutations to affect multiple traits. Since pleiotropic genes are involved in many traits and function in multiple tissues, they are thought to impose genetic constraints on evolution. Changes in these genes may be less tolerated since altering one gene can simultaneously affect several traits, potentially leading to negative effects. The consequences of pleiotropy on evolution have been studied under several theoretical frameworks ranging from Fisher’s geometric model, which assumes universal pleiotropy ^17^, to the seminal work by Alan Orr who showed that as the degree of pleiotropy increases, the rate of adaptation dramatic decreases ^18^. This result gave rise to the idea of a “cost of complexity”, implying that complex animals should have lower rates of adaptation ^18^. Although several modelling approaches have supported this conclusion ^19,20^, the fact that complex organisms have evolved and continue to do so appears to contradict it. The apparent conflict may not stem from the models themselves but rather from their underlying assumption of universal pleiotropy ^21^. Empirical studies in diverse organisms including *Saccharomyces cerevisiae* ^22,23^, *Arabidopsis thaliana* ^24^, *C. elegans* ^25^, threespine stickleback ^26^, mice ^27,28^, and humans ^29^ reveal that the distribution of pleiotropy in the genome is approximately L shaped. These findings suggest that pleiotropy exists as a spectrum, rather than at a fixed level. This raises an important question: If pleiotropy exists as a spectrum, does it always hinder adaptation, or can it sometimes facilitate it?

In this study, we use a data-driven unsupervised approach to identify instances of repeated molecular convergence in protein-coding orthologs across 143 teleost fish species. Teleosts fishes represent the most specious group of vertebrates on Earth, encompassing a wide array of phenotypes, morphologies, habitats, and life histories ^30,31^. This diversity positions teleosts as an ideal group for exploring the broad evolutionary processes giving rise to species diversity and richness. Furthermore, the availability of high-quality genomes for numerous species, coupled with single-cell sequencing and functional data from zebrafish (*Danio rerio*), offers an excellent opportunity to investigate the potential effects variations in coding sequence might have on tissue functionality and animal phenotypes ^32^.

## Results

### Data collection and orthology analysis

We selected teleost fish species that had chromosomal-level annotated genomes from the National Center for Biotechnology Information (NCBI) ^33^ and Ensembl genome databases ^34^. Our final list of genomes comprised 143 species from 40 orders (most sampled were *Perciformes* with 17 species) covering 85 families (most sampled were *Salmonidae* with 10 species) (Fig 1a, Table S1). We use the spotted gar, *Lepisosteus oculatus*, a non-teleost that did not undergo the teleost-specific whole genome duplication ^35^ as our outgroup. Using OrthoFinder ^36^ we estimated orthologous relationships across species. Sets of orthologous genes are then placed in orthgroups. The analysis resulted in 40,940 orthogroups containing over 4 million protein-coding orthologs. 1723 orthogroups contained all the species. To ensure that we focus on genes which reflect the evolution of teleost fish diversity, we selected orthogroups with at least one species from each of the 40 orders in our dataset. Additionally, to keep computational times reasonable we restricted our analysis to orthogroups containing a maximum of 1500 genes. This resulted in a final dataset of 9273 orthogroups with around 2 million orthologous genes (Dataset S1 in online repository), i.e. more than half of all the genes in the original 40,940 orthogroups. Due to this filtering we focus on a more conserved set of genes across taxa. While lineage specific genes can experience different evolutionary trajectories, leading to the evolution of specific (or even the same) adaptations in multiple lineages, the focus of our study is repeated evolution using the same genetic building blocks. We annotated and functionally characterised the excluded orthogroups, and found that they were mostly associated with immune processes and ion transport, whereas the gene sets we used in our analysis were involved in many diverse processes (Fig S1 and S6). Although processes related to ion transport and immunity are important, our sample encompassed a diverse array of processes, and provided a powerful window into teleost evolution.

**Fig 1:**
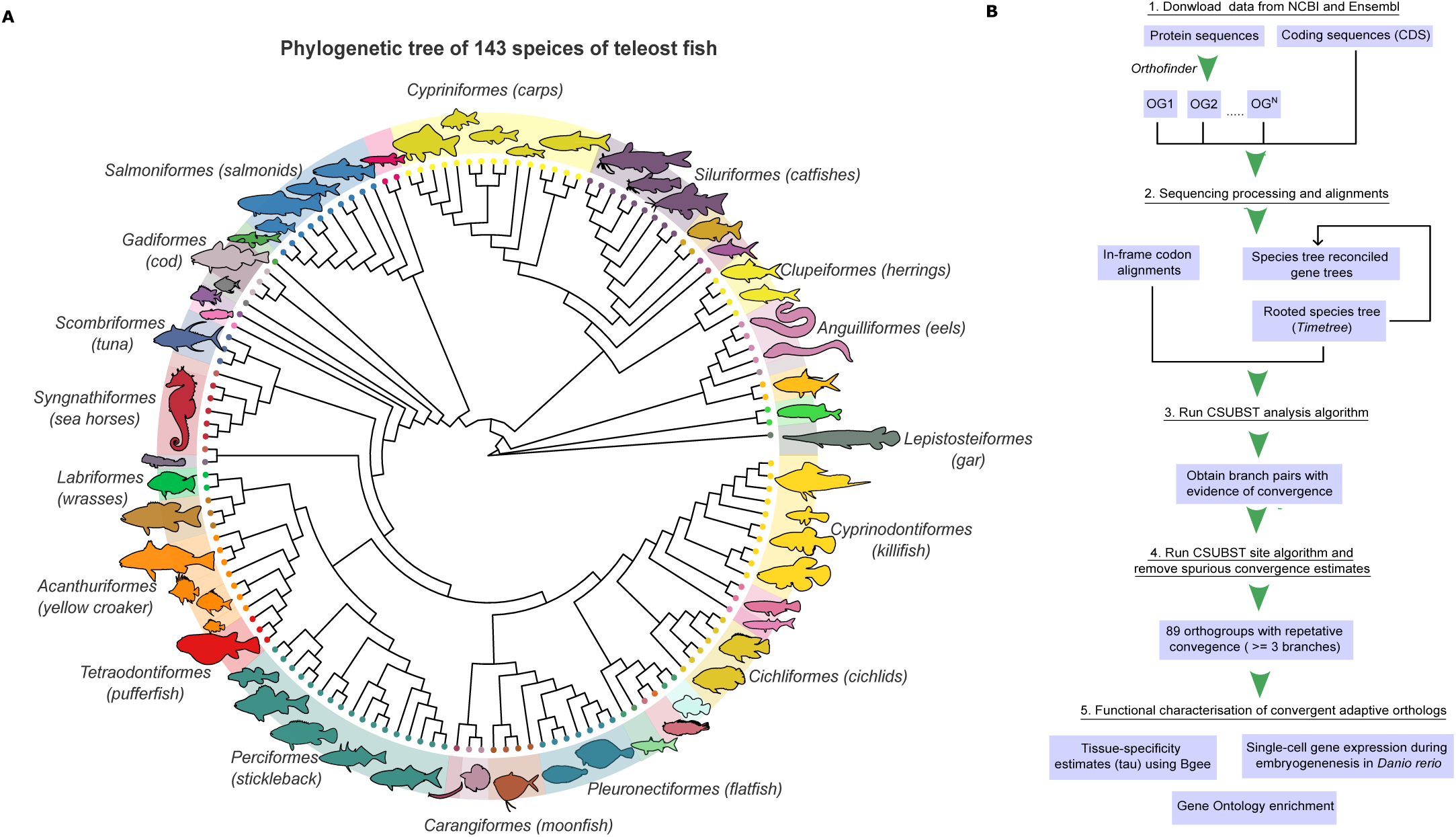
Phylogenetic tree of species, and taxonomic distribution of convergence. (A) Phylogenetic tree of 143 teleost fish species used in the study. The silhouettes are not to scale and are representative of species in each order. (B) Schematic describing the workflow of this study.

### How many genes have evolved convergent substitutions in teleosts?

We obtained estimates for molecular convergence using the CSUBST (Combinatorial SUBSTitutions) algorithm introduced by Fukushima and Pollock ^37^. This algorithm allows for exploration of genomic datasets to identify repeated amino acid substitutions without specifying any foreground lineage or any predefined hypothesis regarding which traits can be considered as adaptations. It combines a pairwise exhaustive search across all possible branch combinations on a phylogeny, followed by a heuristic step that examines branch combinations in more than two independent lineages ^37^. CSUBST draws inspiration from the widely used *dN*/*dS* ratio (parameterised as *ω*) which measures the rate of protein evolution as a ratio between non-synonymous and synonymous substitution rates ^38^. Fukushima and Pollock ^37^ developed a similar metric, *ω*_*c*_, which applies to substitutions repeatedly occurring on combinations of separate phylogenetic branches. This metric is designed to assess the relative rates of convergent evolution by comparing the rates of non-synonymous convergence and synonymous convergence across a range of distinct phylogenetic branches. The rationale behind this approach is that truly adaptive functional changes to proteins are most frequently caused by non-synonymous codon substitutions that alter the amino acid sequence. If distinct (distantly related) lineages exhibit the same amino acid change at the same position this is a strong signal for adaptive evolution as the emergence of such repeated substitution by neutral evolution alone is extremely low. CSUBST generates estimates for various types of combinatorial substitutions. These include divergent changes, characterised by substitutions from any ancestral amino acid to a different extant amino acid, and convergent changes, which involve substitutions from any ancestral amino acid to a specific amino acid occurring at the same position across multiple lineages. Our study specifically focused on convergent changes at the coding sequence level and not at the gene or genomic features levels.

We estimated ωC for all independent branch pairs across 9273 orthogroups and classified branch pairings that jointly satisfied minimum thresholds of 5 for *ω*_*c*_ and for observed non-synonymous convergence (OCN) (*ω*_*c*_ ≥ 5 & OCN ≥ 5) as evidence for convergence. These thresholds were based on validations in the original paper ^37^. We only focussed on convergence that was observed in three or more branch combinations (Table 1). Next, using the CSUBST *site* function we mapped convergent amino acid substitutions on predicted protein structures. Following a final post-processing step to remove certain instances of spurious convergence (Fig S2) (see methods), the final output comprised 89 orthogroups (274 genes), each showing evidence for repetitive convergence in multiple independent branches across teleosts (Table 1). We checked for any potential bias of the OCN metric to estimate convergence for longer genes or orthogroups containing more genes simply because the sequence space is larger. Using a Pearson correlation test, we found no significant relationship between gene number or gene length and the OCN metric (Fig S3). We also compared the distribution of gene copy numbers within the convergent orthogroups and a background of randomly selected non-convergent orthogroups. Interestingly, we found that on average the convergent orthogroups had higher copy number than random sets of orthogroups (1000 bootstrap of one-sided Kolmogorov-Smirnov test; average Benjamini-Hochberg corrected P = 0.002) (Fig 2A).

**Fig 2:**
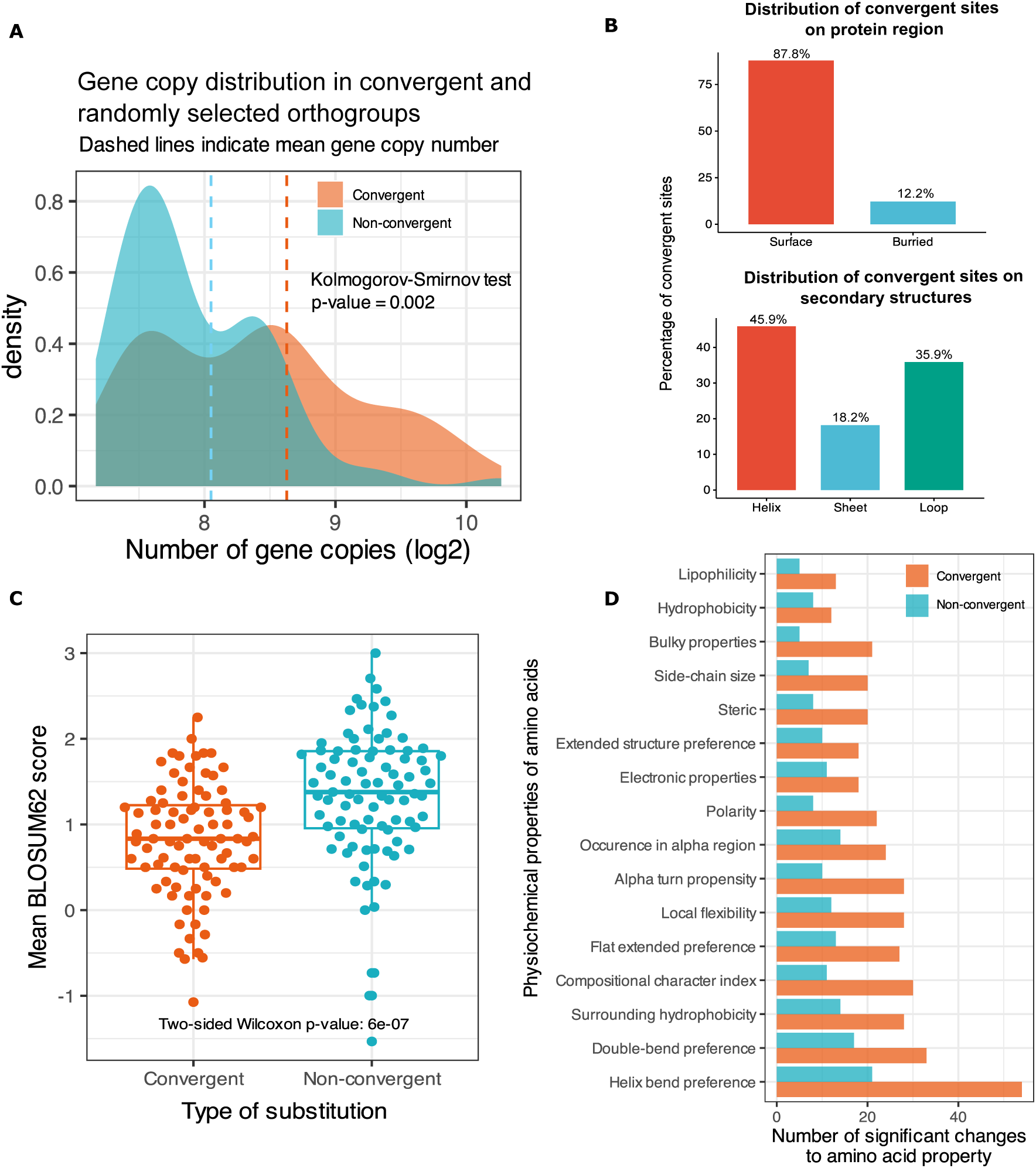
Gene copy number of convergent orthogroups and changes to physicochemical properties following amino acid substitutions. (A) The convergent orthogroups had a higher distribution of genes with high copy number as compared to the background of randomly selected genes. (B) A majority of the convergent sites were on the external surface of the protein, with most of them located on the helices of proteins. (C) Difference in mean BLOSUM62 score during a change from an ancestral amino acid to extant amino acid for the two types of protein substitutions: convergent and non-convergent. The plot shows that the convergent substitutions on an average had lower BLOSUM62 scores as compared to other, non-convergent substitutions occurring within each protein. (D) Comparison of changes in physicochemical property on ancestral and extant amino acids for convergent and non-convergent substitutions. The panel shows that the convergent substitutions had a higher number of statistically significant changes to amino acid properties as compared to the non-convergent substitutions.

**Table 1:**
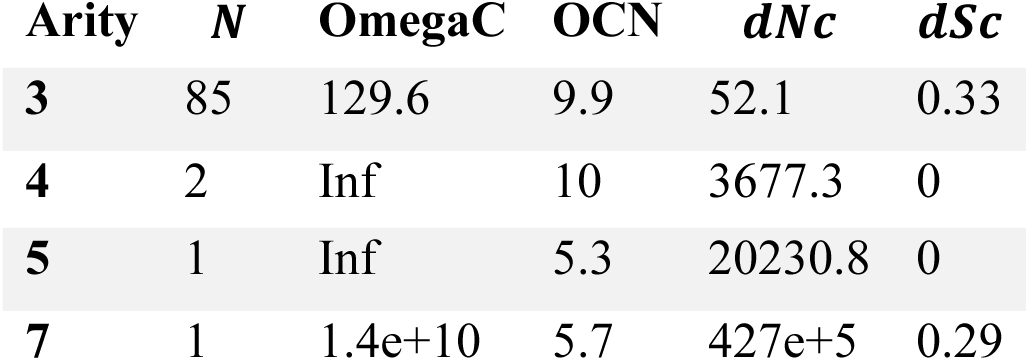
Convergence statistics obtained from CSUBST. Arity is the number of combinatorial branches for which convergence was found. *N* is the number of orthogroups per arity with evidence of convergence. OmegaC median (*ω*_*c*_) value. *dNc* is the median value for the number of non-synonymous convergent substitutions. *dSc* is the mdeian value for the number of synonymous convergent subsitutions.

| Arity | *N* | OmegaC | OCN | *dNc* | *dSc* |
| --- | --- | --- | --- | --- | --- |
| 3 | 85 | 129.6 | 9.9 | 52.1 | 0.33 |
| 4 | 2 | Inf | 10 | 3677.3 | 0 |
| 5 | 1 | Inf | 5.3 | 20230.8 | 0 |
| 7 | 1 | 1.4e+10 | 5.7 | 427e+5 | 0.29 |

Most convergence involved three branches, with a few cases involving four, five, or seven branches (Table 1). The orders *Perciformes*, *Eupercaria*, and *Cyprinodontiformes* had the most convergent genes, as did the families *Cichlidae*, *Percidae*, and *Tetraodontidae* (Fig S4). The taxonomic distribution of convergent genes was different from that simply expected by sampling (two-sample Kolmogorov-Smirnov test P < 0.05) (Fig S4). In terms of ecological characteristics (based on available data from FishBase), the highest proportion of convergent genes were estimated in tropical sea fish (Fig S5). However, we could not infer any overall significant relationship between convergence in a specific orthogroup and ecological features such as climate, ecosystem type, or food items. This was likely due to the low granularity of the categories.

### Estimating functional consequence of amino acid substitutions

Mapping of convergent substitutions onto the protein structure revealed that most convergent sites were located on the surface residues of the protein, potentially enabling interaction with other proteins (Fig 2B). However, convergent sites were not significantly overrepresented on the surface residues (chi-square test P = 0.941). This implies that although most convergent sites were on the protein surface, so were most other sites. In terms of secondary structure, most of the convergent sites were located on helices, but were not overrepresented (chi-square test p-value = 0.567). The heterogeneity of the protein structures resolution likely reduces the statistical power of these comparisons (see methods).

To get an estimate of the function consequences of the convergent substitutions, we compared the average BLOSUM62 scores of the convergent and non-convergent substitutions occurring within each of the proteins with convergent substitutions. We found a significant difference (two-sided Wilcoxon rank sum test P = 6×10^-7^) in BLOSUM62 scores between convergent and non-convergent substitutions (Fig 2C). On an average the convergent substitutions had lower scores implying that these changes are rare in evolution, and have a higher probability to alter protein function. Next, we compared the changes in physicochemical properties of amino acid residues resulting from convergent substitution and non-convergent substitutions within each protein. We compared 17 different amino acid properties and found that 95% of the convergent orthogroups (85 out of 89) had significant changes (two-sided Wilcoxon rank sum test Benjamini-Hochberg corrected P < 0.05) in at least one of the 17 physicochemical properties analysed (Fig 2D). The number of significant changes was higher for the convergent substitutions versus the non-convergent ones. The most frequent convergent changes affected helix bend preferences, which aligns with the observation of their higher distribution on protein helices. However, the highest proportion of significant convergent change were in bulky properties, side-chain size, and alpha turn propensity. The lowest proportions were in occurrence in alpha region, electronic properties, and hydrophobicity. A full description of the properties and their potential effects can be found in Table S2. We also provide several examples of how alterations to amino acid properties can affect various aspects of protein function and stability in supplementary text 1.

### Potential cases of site-specific functions

In cases where it was possible to identify site-specific functions, we found convergent substitutions at critical functional sites. Although specific experiments are needed to validate their function, we can make several predictions based on empirical evidence from previous studies.

For instance, in embryonic haemoglobin alpha 5 protein (*hbae5*; OG0000107), we observed both convergent and divergent substitutions located around the protoporphyrin IX containing Fe ligand binding site (Fig 3A). Changes in the haemoglobin protein, particularly alterations at the binding site of the Fe ligand have been linked to increased tolerance to acute hypoxia in vertebrates ^39,40^. Since we also observed substitutions located close to the Fe ligand binding site, they might impart similar increased tolerance to hypoxic conditions. The convergent substitutions were observed in *Electrophorus electricus* (electric eel), *Larimichthys crocea* (yellow croaker), and *Oreochromis niloticus* (Nile tilapia). Electric eel and Nile tilapia can both resort to air-breathing during times of hypoxia ^41,42^, while the yellow croaker, which is often found in muddy estuaries and coastal waters, has evolved protective, haemoglobin-containing mucous that aids in oxygen transport during hypoxia and air exposure ^43^. The convergent substitutions in *Hbae5* could be linked to these adaptations.

**Fig 3:**
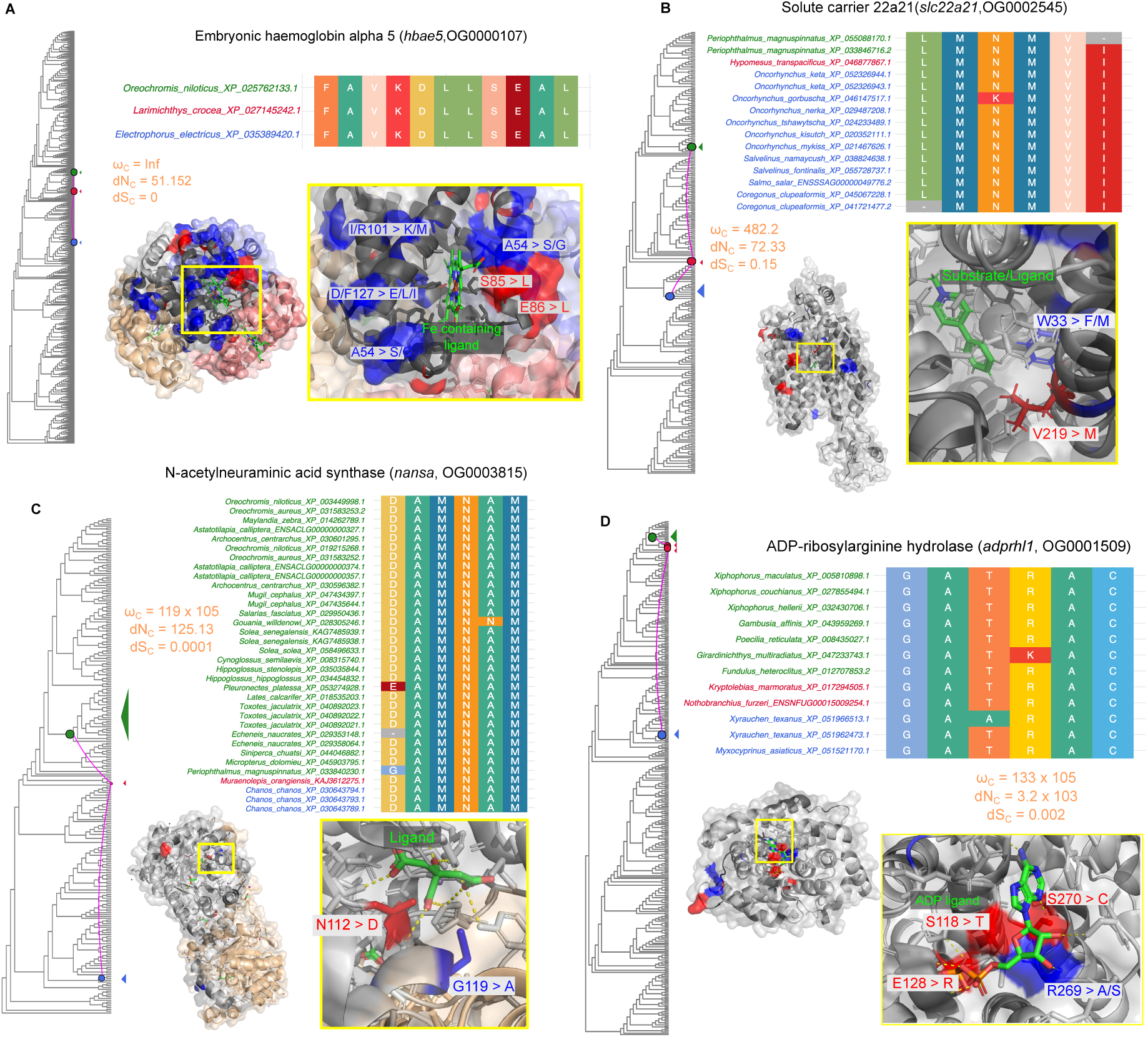
Potential cases of site specific functions. The red sites represent convergent sites while the blue represent divergent sites. The amino acid letter and site followed by the arrow represent the change in sequence at that particular position on the protein structure. For example, in *hbae5*, S85>L means a serine at position 85 on the protein structure changed to a leucine. These sites are localised in important ligand (green) binding sites. The phylogeny is a species-reconciled gene tree. The coloured dots represent the branches in which convergence was found. The species names and colours correspond to the extant species descended from these branches. The alignment shows each of the aligned sites in the species with convergent substitutions. These sites are different from those in other species (full alignment in Dataset S8). Convergence metrics for each branch combination are also shown.

We also detected convergent substitutions within the interior of the channel of the barrel-like structure of the solute-carrier protein *slc22a21* (OG0002545), near the 1-methyl-4-phenylpyridinium ligand binding site (Fig 3B). Given the role of *slc22a21* in maintaining the homeostasis of various organic ions, these substitutions could impact processes related to the absorption, distribution, metabolism, and excretion of biomolecules ^44^. Members of the *Salmonidae* family, as well as *Periophthalmus magnuspinnatus* (mudskippers), and *Hypomesus transpacificus* (Delta smelt), which harbour the substitutions, all experience varying levels of salinity throughout their life cycles ^45–47^. The convergent substitutions could have helped these species adapt to changes in salinity.

The orthogroup OG0003815 encodes N-acetylneuraminic acid synthase (*nansa*). Multiple lines of evidence suggest that *nansa* plays a crucial role in skeletal development, particularly in head formation. Knockdown of *nansa* in zebrafish leads to abnormal skeletal development, while microinjection of *nansa* morpholino into zebrafish embryos results in small head size and developmental anomalies. Additionally, mutations in the human ortholog NANS, which alter its enzymatic activity, have been linked to skeletal dysplasia ^48^. These results support the gene’s essential role in skeletal formation, particularly in the development of the head. The convergent substitutions are present at multiple sites on the protein surface, including a substitution within the protein’s interior at a functional ligand-binding site (Fig 3C). The species within this orthogroup display different body shapes, ranging from the fusiform body shapes of *Orechromis niloticus*, *Astatotilapia calliptera* (eastern river bream), and *Toxotes jaculatrix* (banded archerfish), to the elongated (somewhat anguiliform) body shapes of *Salaris fasciatus* (jewelled blenny), *Muraenoplis orangiensis* (Patagonian morray cod), and *Gouania willdenowi* (blunt-snouted clingfish), as well as the depressiform body shapes of *Soela solea* (common sole) and *Hippoglossus Stenolepis* (Pacific halibut). Interestingly, the remora, which has evolved a flat oval sucking disk (modified dorsal fin) used to attach to larger marine animals like sharks, also harbours substitutions in the *nansa* gene. The occurrence of convergent substitutions appearing in organisms with divergent phenotypes has been observed in marine mammals as well; for instance, substitutions in the *S100A9* and *MGP* genes were linked to both high and low bone density occurring in shallow diving manatees and deep diving dolphins respectively ^49^. Therefore, the convergent substitutions in the *nansa* gene likely influenced the sialylation of glycoproteins which altered downstream pathways affecting head and skeletal morphology.

Finally, ADP-ribosylarginine hydrolase (*adprhl1*; OG0001509) exhibited convergent substitutions at the adenosine-5’-diphosphate ligand site (Fig 3D). This active site is important for the post-translational modifications of proteins ^50^. Although the ADP-ribosylarginine hydrolase gene family affects multiple processes, its link to tumorigenesis might be particularly relevant for the *Xiphophorus* (platyfish) species which readily form skin melanomas ^50,51^.

### Which biological processes are the convergent genes involved in?

We performed a Gene Ontology (GO) enrichment to get a general overview of the biological processes associated with these convergent genes. The analysis revealed a significant enrichment (Benjamini-Hochberg corrected P < 0.05) for terms predominantly linked to biomolecule metabolism, response to hormones or stimuli such as heat, and processes of embryonic development and tissue morphogenesis (Fig S6). Interestingly, we also identified enriched GO terms related to xenobiotic-processing (GO:0071466-response to xenobiotic stimulus), gas transport (GO:0015671-oxygen transport), and sensory system development (GO:0060042-retinal morphogenesis in camera-type eye) which were reported to be frequent targets of adaptation to new environments (Fig 7A) ^16^.

To better resolve the function of these convergent genes, particularly in the context of development and tissue formation, we carried out a comprehensive analysis of gene expression across species and developmental transitions. Using gene expression data from the Bgee database ^52^, we calculated tissue specificity of the convergent orthologs in eleven fish species (see methods) and across eight different tissue types (brain, eye, heart, liver, muscle, ovary, skin, and testis). Approximately one-third (28 out of 89) of the convergent genes were tissue-specific, with the highest number in the liver (Fig S7). Overall, we observed that the proportion of convergent genes that are tissue specific is no different from the proportion of tissue specific genes in our dataset as a whole (Two-sided Wilcoxon signed rank test P = 1), suggesting that the convergent genes are not more likely to be tissue-specific. However, on a tissue-by-tissue basis we found a significant difference in the convergent/non-convergent ratio between tissue-specific and non-tissue-specific genes only for the liver and not for other tissues (Fisher’s exact test P < 0.05); in other words, our convergent gene set had a higher proportion of liver-specific genes than expected by chance. Of note, our estimates of tissue specificity were limited to adult fish.

To gain insight into the expression dynamics of the convergent orthologs during development, we analysed single-cell RNA sequencing data for embryonic development in zebrafish (*Danio rerio*) from the Zebrahub atlas ^32^. We identified three groups of convergent genes based on their expression patterns. First, genes with ubiquitous expression throughout embryonic development and across multiple cell types (Fig 4). These included genes such as *adam17a* (ADAM metallopeptidase 17a; OG0003642), *eef2b* (eukaryotic translation elongation factor 2b; OG0001781) and *eef1a1|1* (eukaryotic translation elongation factor 1 alpha 1; OG0000135) (complete list in Table S3). Involved in the Notch signalling pathway, *adam17a* showed stable expression in the early stages at 10hpf (hours post-fertilisation), 16hpf, and 2dpf (days post-fertilisation) but declined at 10dpf. Both *eef2b* and *eef1a1|1* showed high expression levels across development, with a slight reduction in eye-related cell types at 10dpf.

**Fig 4:**
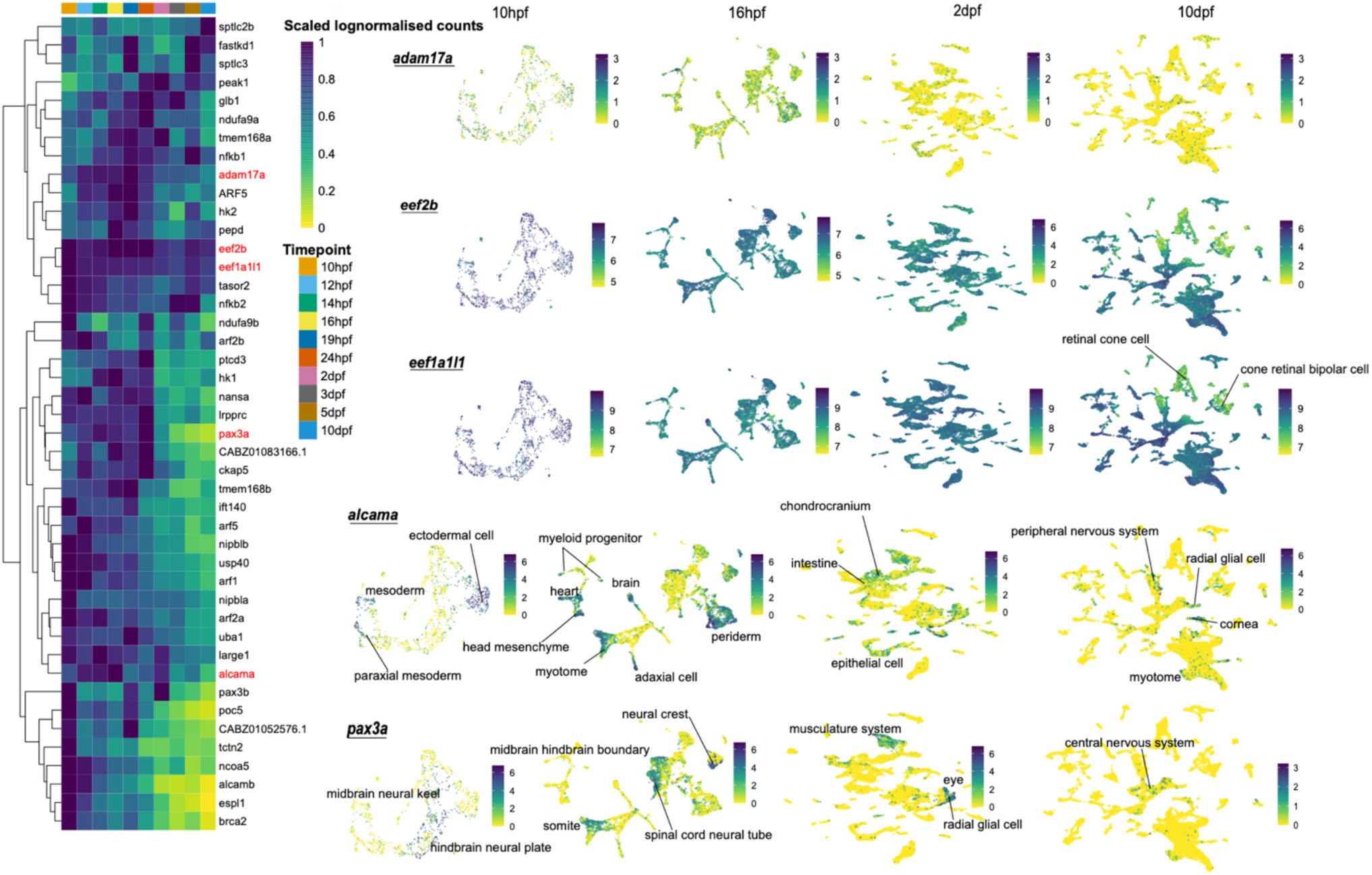
Convergent genes expressed in multiple cell types throughout embryonic development in zebrafish. Single-cell RNA sequencing shows that the convergent genes expressed throughout embryonic development can be divided into two groups. The heatmap shows expression of genes throughout zebrafish embryonic development while the UMAPs show expression of across cell types at specific time intervals. The genes in red on the heatmap correspond to genes in the UPAMs. Genes like *adam17a*, *eef2b*, and *eef1a|1* have expression in almost all cell types. While genes like *alcama* and *pax3a* show more restricted expression in specific cell types. A key feature of both these groups is that the genes are expressed in multiple tissue systems (e.g. heart, musculature, nervous system).The color bar in the UMAP is log-normalised counts.

The second group comprised genes expressed throughout embryonic development but in specific cell types. Examples include *alcam-a* (activated leukocyte cell adhesion molecule a; OG0001823) and *pax3a* (paired box 3a; OG0000460) (Fig 4). *Alcam-a* was highly expressed in cell types related to lens placode, neural crest, radial glial cell, and cornea (Fig 4 10dpf) ^53,54^. With high expression in cell types related to somites, spinal cord neural tubes, radial glial cells, neural crest cells, and the central nervous system, the expression of *pax3a* aligned with its known role in eye development and pigmentation ^55^.

The third group of genes were predominantly expressed late in embryonic development (Fig 5). Genes in this group, such as *slc22a21* (OG0002545) and *hbae5* (OG0000107), were predominantly expressed between 24hpf and 10dpf (Fig 4). *Slc22a21* was associated with the excretory and vascular systems, while *hbae5* was restricted to hematopoietic cells. Other genes, such as *postnb* (periostin, osteoblast specific factor b; OG0002859) and *gstt1b* (glutathione S-transferase theta b, OG0004038), have roles in the formation of the extracellular matrix of the skin ^56^ and liver detoxification, respectively.

**Fig 5:**
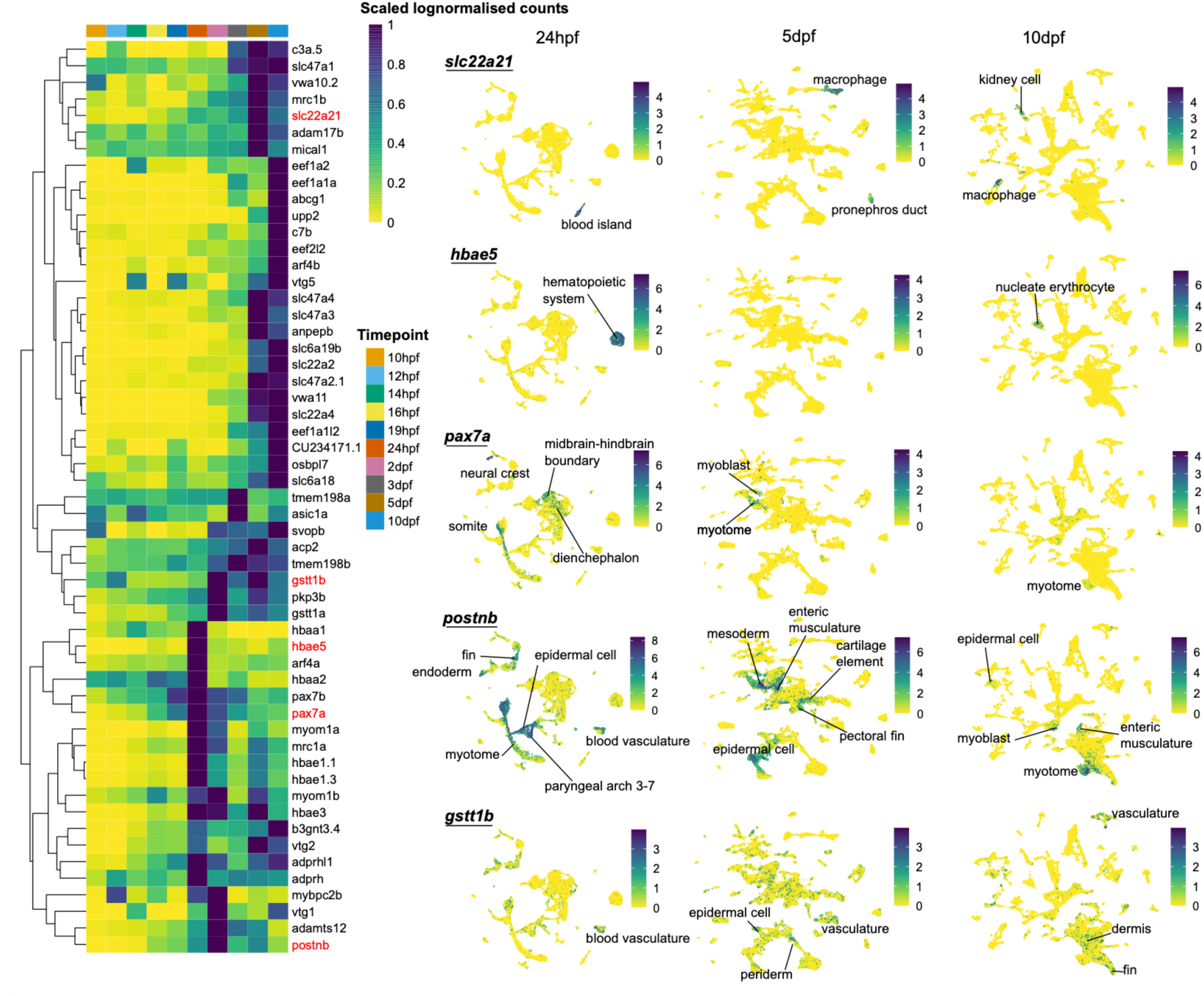
Convergent genes expressed in multiple cell types at specific stages of embryonic development in zebrafish. In contrast to the previous plot, these sets of convergent genes are highly restricted ontogenic expression patterns, being expressed only at specific timepoints. These genes are also expressed in multiple different cell types. The genes in red on the heatmap correspond to genes in the UPAMs.The color bar in the UMAP is log-normalised counts.

### Pleiotropy estimates

The finding that only one-third of convergent genes were tissue-specific, while being involved in diverse processes, and expressed across multiple cell types, suggests that they may be pleiotropic. Directly measuring pleiotropy is challenging due to difficulties in defining relevant traits and determining the appropriate level of measurement ^57^. Given these complexities, multiple lines of evidence are necessary to accurately assess the degree of pleiotropy. Here we combine such multiple lines of evidence to get estimates of biological pleiotropy for our convergent genes relative to background levels. We measured the direct functional role of a gene, determined by its functional diversity, its expression across different tissues, and the number of phenotypic traits affected by genetic perturbations ^57^.

We first analysed the semantic similarities of GO terms between convergent orthologs and a background set of orthologs. We expect more pleiotropic genes to be annotated to more diverse GO terms, and thus have an overall lower semantic similarity among their terms. A critical requirement for accurate semantic similarity analyses are well annotated GO terms, based on both experimental and *in silico* evidence ^58^. Therefore, we carried out this analysis with the *Danio rerio* orthologs. Based on semantic similarity we could classify the convergent orthologs into ten major categories of biological processes (Fig 6A). Some genes, like *alcama*, were annotated across multiple categories—including development, transport, and the immune system—indicating broad functional roles. Others, like *nipbla*, had multiple annotations within a single category, such as development, where it is involved in the formation of the heart, digestive tract, and fins. We found that GO terms of the convergent orthologs, on average, exhibited lower semantic similarity than the background (one-sided Kolmogorov-Smirnov test P < 10^-16^) (Fig 6B). This suggests that they are associated with a diverse range of biological processes ^58,59^, supportive of pleiotropy of the convergent genes. This pattern was consistent across GO terms categories (Fig 6B).

**Fig 6:**
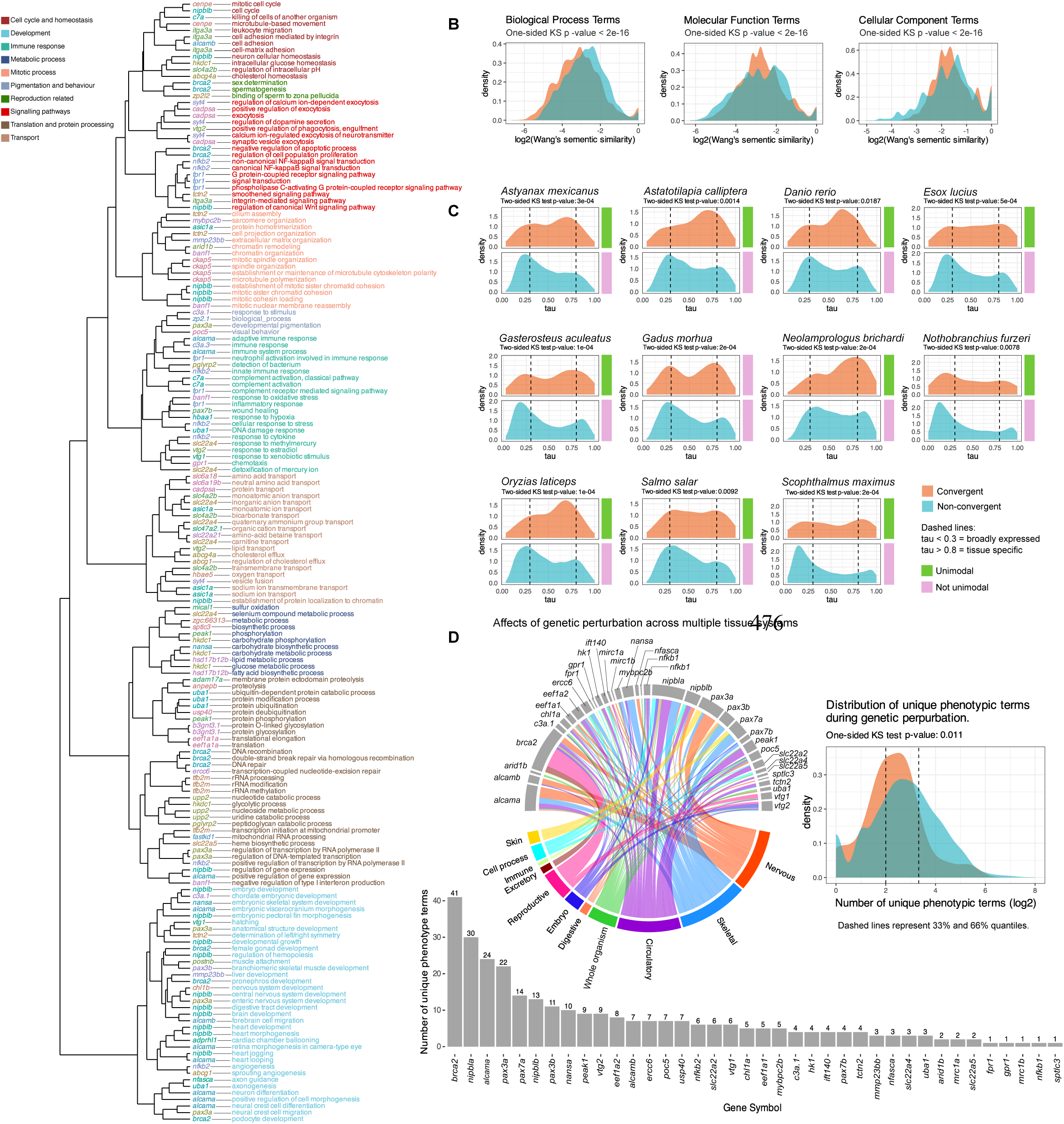
Multiple lines of evidence suggest the convergent genes have intermediate degrees of pleiotropy. (A) Dendrogram of convergent *Danio rerio* orthologs based on GO semantic similarity. The dendrogram shows the different genes and their associated Biological Process GO descriptions, categorised into ten different clusters. Each gene is denoted with a unique colour. Full plots along with plots for other GO categories can be found in Dataset S12. (B) Distribution plots showing that the GO terms of the convergent genes tend to have lower semantic similarity scores as compared to a background set of random genes. (C) The convergent genes had intermediate levels of tissue specificity, i.e most of the genes had tau values between broadly expressed (tau < 0.3) and tissue specific (tau > 0.8) ^66^. (D) Using data from ZFIN, this panel examines the unique phenotypic terms associated with genetic perturbation of convergent orthologs in *Danio rerio.* The histograms show the number of unique phenotypic terms associated with genetic perturbation in *Danio rerio.* These data are obtained from various studies and show that the convergent orthogroups can have multiple phenotypic effects. The chord diagram shows that for some genes the perturbation phenotypes occur across multiple tissue systems (e.g. *alcama*, *pax3a*). The length of the grey bars correspond to the frequency values on the histogram. Data used to construct the chord diagram can be found in Table S5. The density plot shows that compared to the background level or all genes with phenotypic information in ZFIN, the convergent genes affect an intermediate number of phenotypes.

Next, we compared the distribution of tissue specificity of the convergent orthologs to all other genes in each of the eleven species of fish from Bgee. In contrast to the previous analysis where we checked whether convergent genes were more tissue specific than other genes in the genome, this analysis compares the distributions of the values of the tissues-specificity index between convergent genes and all other genes. The distribution of tissue specificity of the convergent genes was significantly different compared to the background (two-sided Kolmogorov-Smirnov test P < 0.05), with the convergent genes having higher frequency at intermediate values of tissue specificity (Fig 6C). Additionally, we used the Silverman ^60^ and Hartigan ^61^ tests to assess the unimodality of tissue specificity of the convergent gene sets. The results indicated that the tissue-specificity of convergent genes in ten out of the eleven species follows a unimodal distribution, in contrast to the bimodal distribution observed in the background set (H_0_: unimodal; test P > 0.05). At the single cell level, we also observed that convergent genes were expressed in an intermediate number of cell types rather than at the extremes of the distribution (Fig S8). However, we did not find any evidence of unimodality using either test, likely due to the sparse nature of the single-cell expression matrix.

Lastly, we collected data from the Zebrafish International Network (ZFIN) database ^62^ to examine the distribution of unique phenotypic terms associated with genetic perturbation (knockdowns, mutation screens, and morpholino experiments) in *Danio rerio*. The ZFIN database provides a wide array of expertly curated cross referenced genetic and genomics data that is routinely used to study gene functions ^62^. For the convergent genes with data in ZFIN, we observed that genetic perturbation can affect several structures across multiple tissue systems (Table S4)(Fig 6D). Compared to the background of all genes with available phenotypic data, the convergent genes had a higher distribution at intermediate levels of affected phenotypic terms, significantly different from the background (one-sided Kolmogorov-Smirnov test P = 0.011) (Fig 6). Collectively, these findings classify the convergent genes as exhibiting intermediate levels of pleiotropy, supporting previous predictions that moderate pleiotropy facilitates adaptation ^63–65^.

### Linking convergence, pleiotropy, and adaptation to ecological shits

To reconcile our observation of multiple convergent substitutions in genes with intermediate levels of pleiotropy across distantly related fish species we introduce the pleiotropic release (PR) model (Fig 7A). Consider a multidimensional adaptive landscape where ancestral species initially reside at distinct adaptive peaks. An ecological shift introduces a new adaptive optimum. Both species now reside at low fitness regions and must adapt by moving toward the new peak. There are two possible evolutionary routes: (i) gradual adaptation via low-effect, low-pleiotropy genes, which drive small incremental phenotypic changes, or (ii) rapid adaptation via pleiotropic genes, which affect multiple traits simultaneously, enabling larger shifts in phenotype. This model predicts that during the initial periods following ecological shifts, adaptation along the lines of pleiotropic genes would be favourable as species can escape low-fitness regions more quickly, and converge on the shared adaptive peak and attain their extant condition.

**Fig 7:**
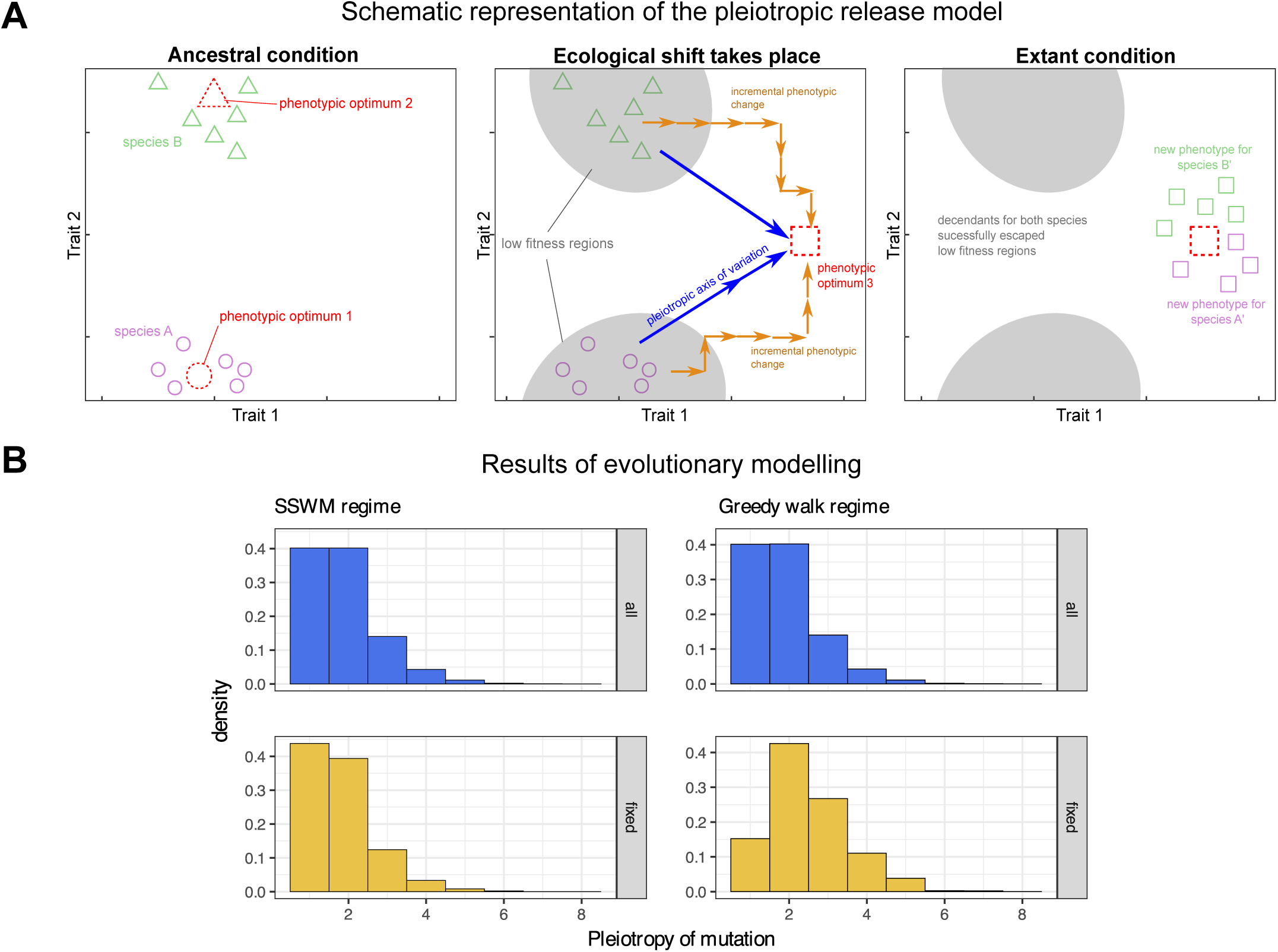
The pleiotropic release model. (A) In the ancestral condition two species occupy separate phenotypic optima in an adaptive landscape. The circle and triangle represent different phenotypic configurations. Following an ecological shift, the optima changes and the species now occupy regions of low fitness (grey). Escape from the low fitness regions can happen through either incremental changes in low, single effect genes (orange) or through large effect pleiotropic genes (blue). Changes along the pleiotropic axis of variation will be favoured as they will enable the species to escape the low fitness regions faster. The optimum can be reached through a few large effect genetic changes, which can lead to convergent amino acids. Overtime, species will evolve a convergent phenotype and occupy the new optima in the extant condition. (B) We tested the predictions of the pleiotropic model by modelling the evolution of populations under two adaptive regimes. In the SSWM regime the frequency of all available and fixed mutations is the same. However, in the greedy regime the frequency of fixed mutations shifts to the left, denoting a higher probability of mutations with intermediate levels of pleiotropy to fix in the population. This suggests that during the initial periods of evolution following an ecological shift, adaptation can occur via the axes of pleiotropic variation.

To test the PR model’s predictions, we use an evolutionary modeling approach based on Fisher’s geometric model (FGM) ^17^. While, various studies have explored adaptation to a constant or moving optima using the FGM ^67,68^, these models assume universal pleiotropy—wherein, every mutation affects all traits that determine the fitness. However, recent studies show that most genes exhibit low pleiotropy, with only a few displaying high pleiotropy, and that mutations in genes affecting more traits tend to have larger per-trait effects ^63,69,70^. We incorporate these empirical trends into our model and track the pleiotropy of genes that acquire mutations during the early stages of adaptation. Using a distribution of pleiotropy derived from empirical data ^63^, we ran adaptive walks using two widely used evolutionary regimes: Strong Selection, Weak Mutation (SSWM) Regime, where mutations are rare but, when they occur, strong selection ensures the fixation of beneficial mutations and the loss of deleterious mutations ^71^. This causes the population to be monomorphic at all times. This regime represents adaptation in an idealised population with low genetic diversity and strong selection. The second regime used is the Greedy Adaptive Walk Regime ^72^, where multiple mutations arise (e.g., due to an increase in population size) and the mutation with the highest fitness advantage fixes in the population. This regime models multiple segregating sites, such as that occurring after ecological shifts. To determine the specificity of our model, we also modelled pleiotropy using two artificial distributions, a reverse L shape where most genes have high pleiotropy, and a uniform distribution where all genes have the same pleiotropy.

The results of the adaptive walks using the empirical distribution showed that in the SSWM regime, the pleiotropy distribution of fixed mutations closely mirrored that of all mutations that arose, with slightly lower mean pleiotropy among fixed mutations, as expected due to limited mutation availability (Fig 7B). In contrast, under the greedy walk regime, the pleiotropy distribution of fixed mutations shifted, favoring mutations with intermediate pleiotropy (Fig 7B). For the reverse L distribution, both regimes had approximately the same distribution of available and fixed mutations (Fig S9). In the uniform distribution, the SSWM regime fixes nearly all available mutations with a slight preference for those with lower pleiotropy (Fig S9). This was expected because, with only a few mutations available, there is no competition between high- and low-pleiotropy mutations. However, since mutations can have either positive or negative fitness effects, lower-pleiotropy mutations, which generally have smaller effects, are more likely to fix. In contrast, the greedy adaptive walk regime shows a strong preference for highly pleiotropic mutations, which was also expected. In this regime, where many mutations are available, those with higher pleiotropy and positive fitness effects tend to fix first. As a result, adaptation in this scenario predominantly occurs through pleiotropic mutations.

Our modeling results suggest that in mutation-rich environments, genes with intermediate pleiotropy play a crucial role in early adaptation to ecological changes, supporting our empirical observations. Furthermore, the pattern of intermediate pleiotropy emerges only when using a real-world empirical pleiotropy distribution, not with artificial distributions, highlighting the specificity and biological relevance of our model.

## Discussion

### Molecular convergence and pleiotropy

Although traditionally considered as a source of genetic constraints ^18^, studies in several model systems as well as natural populations have shown that the idea of pleiotropy constraining adaptation needs to be reconsidered ^57,73^. For example, a vital gene responsible for the adaptation to freshwater habitats in sticklebacks, Ectodysplasin (*Eda*), is highly pleiotropic and responsible for ectodermal appendage formation, sensory system patterning, and schooling behaviour ^74,75^. Intermediate levels of pleiotropy might even enhance adaptive potential by causing greater fitness gain and increasing the fixation of beneficial alleles ^63,76^.

Multiple models have previously studied the effect of pleiotropy on adaptation. For instance, Otto, 2004 showed how adaptation can be constrained and the mean selection coefficient is halved under the assumption of the total pleiotropic effects of an allele being negative ^20^. However, this model focused on alleles that affect few focal traits, in an otherwise well adapted organism. Our modelling approach differs from this, since we assume that populations are far from the optima in each trait dimension following an ecological shift, and we do not impose any conditions on the total pleiotropic effect – these differences can explain why we observed a preference for mutations in genes with intermediate pleiotropy. Another study ^77^, much like ours, combined a genetic architecture with Fisher’s geometric model, however they focused on ‘orientational heterogeneity’ in the effects of mutations, while considering each locus to have the same pleiotropy. Our model differs from this since we sample pleiotropy as a spectrum based on an empirical distribution. Chevin *et. al.* found that the probability of parallel evolution is higher for lower pleiotropy, when all loci have the same pleiotropy ^77^. However, they also predicted that pleiotropy heterogeneity could increase the probability of parallel evolution ^77^.

We can use our model to explain the patterns that we observe in our study. For example, an environmental shift might have resulted in the ancestors of salmonids, mudskippers, and delta smelt to each experience varying levels of salinity. The varying levels of salinity could also impact the available food resources as food webs differ between marine and freshwater habitats. To adapt to the new environmental conditions changes in the *slc22a21* gene might prove particularly beneficial. Along with maintaining the homeostasis of organic ions between tissues and the interfacing body fluids, this gene family can impact tissue systems such as the liver, kidneys, skeletal system, and central nervous system^78^. Additionally, they also facilitate inter-organism communication between the gut microbiome and the other tissue systems^78^. Therefore, adaptive variations in *slc22a21* gene could cumulatively enhance liver detoxification, improve ion reabsorption by the kidneys, and strengthen signalling between the gut microbiome and the central nervous system, all of which would be particularly beneficial during ecological shifts involving changes in salinity and dietary changes as a result of altered nutrient availability. Considering that teleosts occupy diverse niches in different environments, evolution through changes in pleiotropic genes might have been particularly relevant during the initial periods of colonisation into new niches.

It is important to note that the pleiotropic release model does not suggest a lack of cost on multiple traits. Adaptation along the axes of pleiotropic genes would likely have costs, but the relative increase in fitness would outweigh these costs. Our model offers an explanation for how multiple distantly related species can reach the same phenotypic optima by accumulating adaptive variation in genes of intermediate pleiotropy. While further studies are needed to assess the generality of the model across taxa and evolutionary contexts, our results provide a solid foundation for future exploration.

### Evolution using pleiotropic genes

In a recent experimental evolution study in yeast, Grant *et. al. ^79^* showed that large effect adaptive mutations are generally pleiotropic, improving both respiration and fermentation, and tend to occur during the early stages of evolution. Although they observed a general shift from pleiotropic to modular adaptation, they did observe some strongly adaptive yeast clones that continued to improve in both respiration and fermentation causing them to achieve high levels of fitness. The authors also discuss observations in multiple studies which show that when yeast are evolved in a glucose-limited environment, the first adaptive mutations are always in pleiotropic genes that enhance both respiration and fermentation. Therefore, pleiotropic release via evolution along the axis of pleiotropic genes might be a feature of evolution during the initial periods following ecological shifts.

Genetic redundancy brought about by gene or genome duplications, and modularity, are two possible mechanisms that can overcome the adaptation-impeding effects of pleiotropy ^57^. In our analysis, the number of gene copies is generally higher in the convergent orthogroups than in the non-convergent ones (Fig 2B). Additionally, some genes show clear differences in the expression of their paralogs (Fig S10). The presence of multiple copies coupled with the divergence in expression patterns between paralogs could enable partitioning of effects, allowing for variation among paralogs to persist non-adaptively as cryptic divergent variations (hereafter cryptic divergence). Cryptic divergence is the genetic difference between paralogs variations that have negligible or no phenotypic effects, but during periods of environmental shifts, produce heritable phenotypic variation. It is analogous to cryptic variation among alleles ^80^. The developmental impacts of cryptic variation in pleiotropic genes have been experimentally demonstrated in models like *Drosophila melanogaster*, *C. elegans*, *Arabidopsis thaliana*, and zebrafish ^81–84^. *In vivo* studies in natural environments, such as those in sticklebacks and cavefish, have highlighted the critical role of ancestral cryptic variation in facilitating the adaptation to ecological changes ^85,86^. Additionally, experimental evidence indicates that mutations in protein-coding sequences of enzymes can generate cryptic variation which maintains the enzyme’s native function on its ancestral substrate while enhancing its adaptive potential in a novel substrate ^87,88^. Therefore, a change in environment can cause the cryptic variation to manifest, leading to phenotypes with enhanced fitness. While not the focus of this study, it is important to note that along with changes to the coding sequence of a gene, variation in gene regulatory elements can have a strong effect on the evolution of phenotypes across diverse scenarios. For instance, alterations to the enhancer of the *Pitx1* gene has been linked to the loss of pelvic structures in certain populations of stickleback ^89^. Considering that teleosts have experienced multiple rounds of genome duplications, repeated loss and modification of regulatory elements can also have important adaptive roles.

Our study has a few caveats. A major requirement for our study was the need for fully annotated chromosomal level genomes. Most of the teleost fish genomes that fit this criteria were of tropical and subtropical fish species, and comparatively fewer species representing temperate, boreal, and polar fish. Although most species of fish are tropical and subtropical, studying the extent of convergence in other groups would be particularly interesting considering that temperate and boreal fishes have smaller effective population sizes than tropical fish ^90,91^. Increasing the number of temperate fish would provide insight into how their specific demographic features can influence the degree of convergence. For instance, the convergent patterns we observe may require large effective population sizes where selection as opposed to drift in the primary evolutionary force. As a result we may observe less convergence in non-tropical fishes because of their relatively lower population size and stronger influence of genetic drift, particularly in freshwater fish ^92^. Another caveat is the absence of gene expression and genetic perturbation data for the species that exhibit the convergent substitutions, which forced us to use zebrafish as a proxy. It is possible that the convergent genes in these species display a more tissue specific expression, functioning preferentially in their target tissues as compared to the other tissues. This could, in turn, reduce the pleiotropic effect of mutations in these convergent genes, suggesting a coordinated evolution of gene regulation and coding-sequences. It would be particularly interesting to explore the possibility of coevolution of mutational effects at the cis-regulatory and coding sequence level ^21^. This possibility presents an alternative explanation for our findings, which we aim to explore in future studies.

## Data availability

All data comprising sequencers, output files, figures, code, and Datasets are available in the online repository: https://doi.org/10.5281/zenodo.15039717

An electronic report containing output of R code and major analyses in this study can be found at: https://agneeshbarua.github.io/Teleost_convergence/

## Materials and Methods

### Data collection and estimating orthologs

In our study, we obtained chromosomal-level annotated teleost fish genomes from the NCBI genomes ^33^ and Ensembl genome databases ^34^ (Table S1). Using an initial list of species we obtained taxonomic information from FishBase (Froese, R. and D. Pauly. Editors. 2023. www.fishbase.org,) and a time-calibrated, rooted species tree from timetree.org ^93^. We only included species that were present in the species tree and downloaded their protein and coding sequences. Using the protein sequences from a final set of 143 species, we constructed orthologs using OrthoFinder ^36^, resulting in 40,940 orthogroups containing 98.2% of all the genes (Table S4). The final dataset consisted of 9224 orthogroups with around 2 million orthologous genes (Dataset S1 in online repository).

### Processing of the orthologs

Prior to running the CSUBST algorithm, we executed a series of preprocessing steps, closely following the guidelines outlined in the CSUBST wiki (https://github.com/kfuku52/csubst/wiki/Preparing-CSUBST-inputs). First, we used the CDSKIT v0.91 tool (https://github.com/kfuku52/cdskit) to isolate the longest isoform for each gene and make the sequences in-frame. Next, using TRANSeq v6.6.0^94^ we translated the coding sequences into protein sequences followed by alignment using MAFFT v7.481 with the ‘auto’ option ^95^. We then performed a translation alignment using TRANALIGN v6.6.0 ^94^ and trimmed poorly aligned sequences using TRIMAL v1.4.1^96^ with the ‘*ignorestopcodon*’ and ‘*automated1*’ options. Finally, we masked and removed ambiguous sites in the alignment using the ‘*mask*’ and ‘*hammer*’ functions of CDSKIT v0.91.

We used IQ-TREE v2.0.3 ^97^ with the MPF model, and 1000 ultra-fast optimised bootstrapping replicates (‘*bnni*’ option), to construct gene trees. We rooted and reconciled the gene trees with the species trees using a combination of GeneRax v1.1.0 ^98^ and NWKIT v0.10.0 (https://github.com/kfuku52/nwkit).

Detailed scripts and commands utilised throughout these steps can be found within the ‘Scripts’ folder in the online repository. We also provide a conda environment YAML file that users can use to recreate the environment for the current pre-processing step as well as the following analysis step. We ran the convergence analysis using a rooted time-calibrated species tree obtained from timetree.org ^93^.

### Convergence analysis using CSUBST

CSUSBT employs a metric similar to the popular *dN*/*dS* ratio (parameterised as *ω*) which represents the rate of protein evolution as a ratio between non-synonymous and synonymous substitution rates ^38^. Distinct from *ω*, the *ω*_*c*_ parameter captures repeated substitutions across various combinations of phylogenetic branches. These combinations range from a minimum of two branches, indicating pairwise substitutions, to a maximum of ten branches. The software quantifies these combinations as *arity (K),* where *K* = 2 represents pairwise comparisons, and *K* = 3 signifies a three-way branch comparison. By estimating all possible combinations of amino acid substitution (across multiple branch combinations) occurring in a multi-dimensional sequence space, *ω*_*c*_ measures how much proteins in different lineages have moved towards the same evolutionary endpoints in this sequence space. CSUBST reports *ω*_*c*_ statistics for nine types of combinatorial substitution ^37^. We focus primarily on *omegaCany2spe* - denoting combinatorial substitution from any ancestral codon to a specific descendant codon, namely, convergent substitutions - and *OCNany2spe*, which denotes the observed rate for non-synonymous convergence. We ran the CSUSBT algorithm using the ‘CSUBST *analyze*’ command with five threads and conducted an exhaustive search for a max *K* = 10. The output of this command can be found in Dataset S4 in the online repository. Orthologs were categorised based on the presence of convergence across these branch combinations and further analysed using the ‘*CSUBST site*’ command (input files in Dataset S5) to map convergent amino acids onto protein structures. This mapping facilitates the visualisation of convergent amino acid sites, particularly their proximity to functional domains within the protein. We employed ad-hoc filtering to remove spurious convergence estimates (Fig S2). In this post processing step, we identified amino acid substitutions that occurred in very close proximity localised at particular regions of the protein. These are usually caused by misaligned sequences due to splice variants between species. After excluding cases of spurious convergence we identified 89 orthogroups that showed evidence for reliable convergence. Dataset S6 contains the non-spurious convergence results, while Dataset S7 has examples of spurious convergence.

### Analysis on protein structures and amino acid substitutions

The ‘*CSUBST site*’ command obtains protein structures by searching the RCSB PDB database (RCSB.org) ^99^. If no experimentally determined structures are found, CSUBST uses NCBI’s QBLAST ^33^ search against the UNIPROT database ^100^ to obtain a .pdb file from the AlphaFold Protein Structure Database ^101^. As a result, the structures obtained are often heterogeneous, comprising X-ray crystallography structures of protein in complex with a ligand, non-complexed protein in crystal, or computationally estimated structures from AlphaFold. This heterogeneity makes statistical comparisons of protein sites difficult. Nonetheless, we applied a systematic selection method to obtain as objective a comparison as possible. We used Pymol ^102^ version 2.4.5 to systematically classify the residues. For each Pymol session, convergent residues were selected based on their colour (red), surface residues were determined using a cutoff of 1.4 Å^2^ to determine their exposure to the solvent. Residues with a distance less than equal to 4 Å from the ligand were classified as in the vicinity of the ligand. Buried residues were those that did not have access to the surface or internal cavity of the protein. The secondary structures of helices, sheets, and loops are selected directly using the keywords provided in Pymol. For all these selections, a second b-factor parameter > 0 was added to eliminate residues that were present in the sequence but not built into the structure. From this set, we can then cross-referenced the selections to determine the desired list and number of residues. These results are in Table S6. Using the *CSUBST site* output files (Dataset S6) we obtained the ancestral and extant amino acids for both convergent and non-convergent substitutions. We compared the average BLOSUM62 scores between the two types of substitutions using the pwalign R package ^103^. We obtained data for the physicochemical properties of amino acid residues from the Peptide R package ^104^. For each protein we obtained the values of the ancestral amino acid properties for the two types of substitutions and tested for significant changes from the ancestral condition to the extant convergent condition and filtered out the number of significant changes (FDR < 0.05)(Fig 2E). Figures showing the difference in properties for each protein can be found in Dataset S11.

### Analysis of GO terms

We used DeepGOPlus v1.0.15 ^105^ to annotate all the genes (∼4 million) in our dataset with gene ontology (GO) terms. DeepGOPlus integrates a convoluted neural network along with sequence similarity scores of protein sequences to annotate GO terms. In our analysis, we use a prediction threshold of 0.3 ^105^. Using the GOstats and GO.db Bioconductor ^106^ packages we performed GO enrichment for GO terms related to Biological Processes. In this analysis, convergent orthologs constituted the test set, while the remainder served as the background universe. Enriched terms were identified using a BH-false discovery rate cut-off of <0.05. For the semantic similarity analysis we used GO annotation of *Danio rerio* orthologs obtained from GO.db ^106^, and carried out the analysing using the GOSemSim R package ^59^. We computed pairwise semantic similarity between GO terms for two gene sets; the convergent *Danio rerio* orthologs and a random set of orthologs used as background. We used multiple random sets and always obtained the same results. We used the *Wang* measure of semantic similarity using *BMA* aggregation ^59^.

### Bgee data and estimating Tau

To gain deeper insight into the functional relevance of the convergent genes we used data from the Bgee database ^52^ and studied expression across eleven species (*Astyanax mexicanus, Astatotilapia calliptera, Danio rerio, Esox lucius, Gasterosteus aculeatus, Gadus morhua, Neolamprologus brichardi, Nothobranchius furzeri, Oryzias latipes, Salmo salar,* and *Scophthalmus maximus*) and eight tissues. We calculated tissue specificity for the brain, eye, heart, liver, muscle tissue, ovary, and testis since they were the tissues with the most consistent sampling across species. All species had gene expression data for these tissues. To calculate tissue specificity, we used the Tau metric because of its efficiency in recovering biological signals ^66^. We classified a gene as tissue-specific if it had a tau >= 0.8 and its expression in the target tissue was greater than the sum of its expression in other tissues. Since we are comparing tissue specificity across different species, we identified genes that are tissue-specific in the same tissue across all the species sampled. These criteria ensured we captured robust signals for tissue specificity.

### Analysis of single-cell RNA-seq from Zebrahub

We used single-cell RNA-seq (scRNAseq) from the Zebrahub consortium ^32^. This data comprises a high-resolution scRNAseq atlas of zebrafish development achieved by sequencing individual embryos across ten developmental stages ^32^. The dataset contained 120,444 cells and a total of 529 cell clusters which were annotated based on published literature and the ZFIN database ^62^. The entire dataset is available at https://zebrahub.ds.czbiohub.org/. The consortium provides h5ad objects with quality control, clustering, and uniform manifold projections (UMAP) (https://github.com/czbiohub-sf/zebrahub_analysis). We log-normalised this processed data using the scater R package ^107^ and use it for our study. To visualise the expression of the convergent genes across embryonic development we calculated the mean expression of each gene at each time point and constructed a scaled (max scaled to 1) heatmap using the scuttle ^107^ and dittoSeq ^108^ R packages. We visualised heatmaps using the plotReducedDim function from the scater R package ^107^. All UMAP plots with cell type and gene expression at each time point are available in Dataset S9. To estimate distribution of gene expression in cell types we first filtered out low expressed genes (*log counts < 0.5*) and then selected cells that had expression across all the sampled fish for each timepoint. This was done to ensure we obtain a consistent gene expression profile across timepoints. We then calculated the number of unique cell types each gene was expressed in. Using this data we estimated the distribution of cell type expression for the convergent and non-convergent gene sets.

### Evolutionary modelling

We considered a genotype of length *L* with *K* alleles at each site (*L* = 10, *K* = 4). An *n*- dimensional vector (*p̅*) stored the phenotypic traits that determine the fitness, based on Euclidean distance from an environmentally determined optimal phenotypic trait vector (*pₒ̅*). Following Fisher’s geometric model, we employ a Gaussian fitness kernel *F(p̅),* with the parameter *σ* fixing the width of the fitness function i.e. the strength of selection.

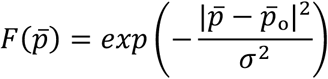

Keeping in mind the recent observation that parallel evolution is driven by SNPs in highly pleiotropic genes ^70^. We only considered point mutations in our model. The number of traits that were affected by this mutation (i.e., the pleiotropy P, 1 < *P* < *n*, of the gene in which the mutation occurs) was sampled from three different distributions: A negative binomial distribution *NB(a, b)* with parameters *a* = 1 and *b* = 10 that were chosen because the distribution mimics the right-skewed empirically observed distribution found in ^63^. A reversed negative binomial distribution, with a left skew generated by subtracting a random variable *r* ∼ *NB(a, b)* from the total number of traits *n*. This was used as a control to understand the effect of the pleiotropy distribution on our results. A uniform pleiotropy distribution *U(1, n)*.

Depending upon the distribution, *P* out of the *n* phenotypic traits were then mutated, with the fitness effect of the mutations sampled from a Normal distribution *N(0,1)* with zero mean and unit variance. We also repeated the analysis with fitness effects of mutations scaled with the pleiotropy of the gene in which the mutation occurs, i.e., each normally distributed random variable was multiplied by the pleiotropy of the gene. We used two different evolutionary regimes: The mutation-limited strong selection weak mutation (SSWM) regime (SSWM), in which each mutation that appears either fixes in the population with a fixation probability,

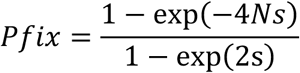

where *s* is the selection coefficient and *N* is the population size. The mutation-surplus greedy adaptive walk regime, in which the fittest of all 1-mutation neighbors of the current genotype fixes in the population. These evolutionary steps are iterated until the global optima is reached. Along the evolutionary simulations, we store the pleiotropy of each mutation that occurs and each mutation that fixes in the population. Comparing the pleiotropy distribution in the former and latter cases allowed us to assess whether early stages of adaptation are driven by mutations in pleiotropic genes. The output of the simulations can be found in Dataset S13.

