## Supplementary material for "Estimates of molecular convergence reveal genes with intermediate pleiotropy underlying adaptive variation across teleost fish": Fig S

An electronic report containing output of R code used in this study can be found at:

https://agneeshbarua.github.io/Teleost_convergence/

All data comprising sequencers, output files, figures, and code are available in the Zenodo data repository: https://doi.org/10.5281/zenodo.15039717

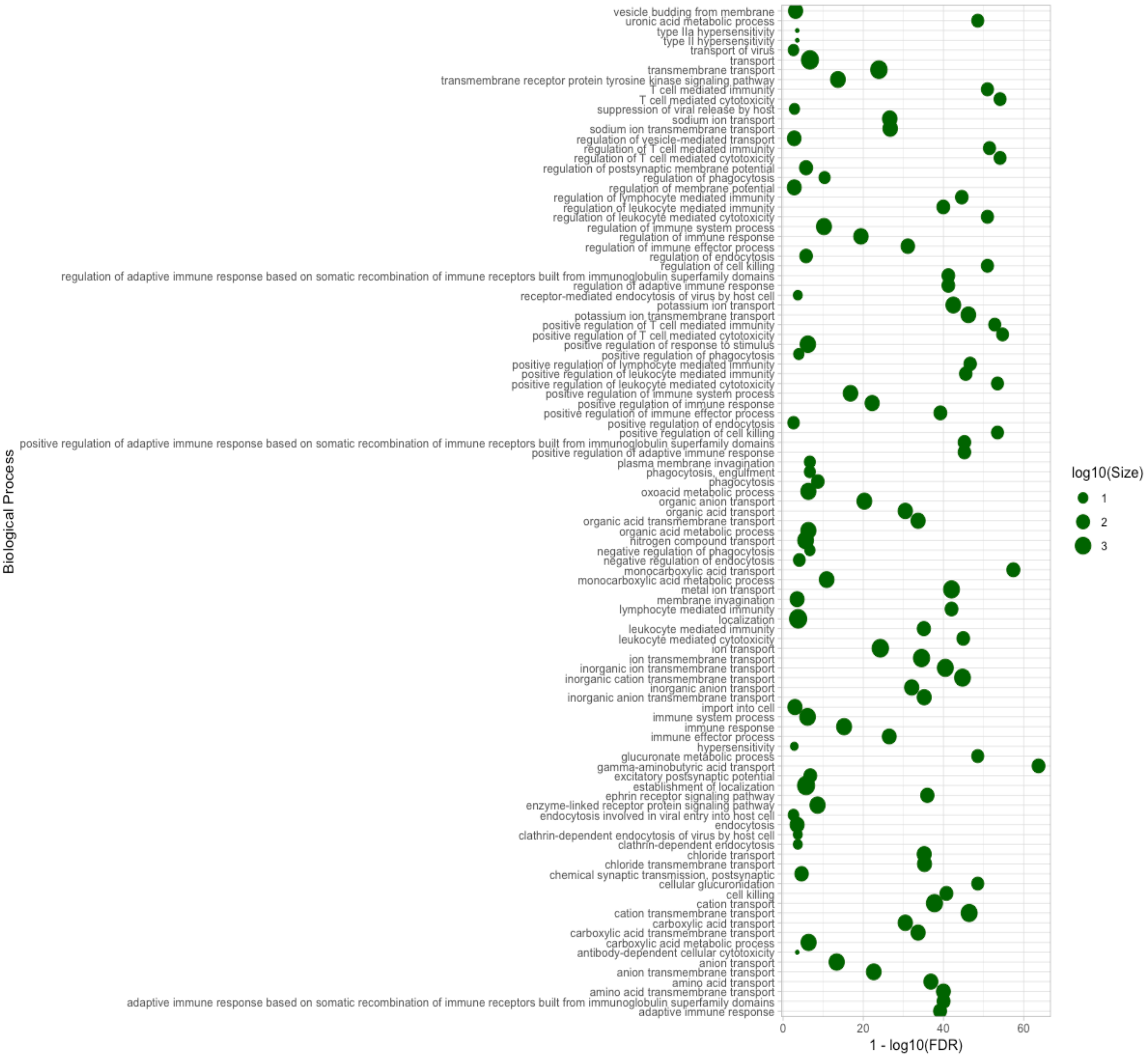


**Fig S1: Gene Ontology (GO) term enrichment of excluded orthogroups.** To keep computational times reasonable we restricted our analysis to orthogroups containing a maximum of 1500 genes. We annotated and functionally characterised the excluded orthogroups and found that they were mostly associated with immune processes, metabolisms, and cell signalling. Although processes related to cell signalling and immunity have substantial functional significance in organisms, our sampled gene sets encompassed a more diverse array of processes (Fig S6).

**A**

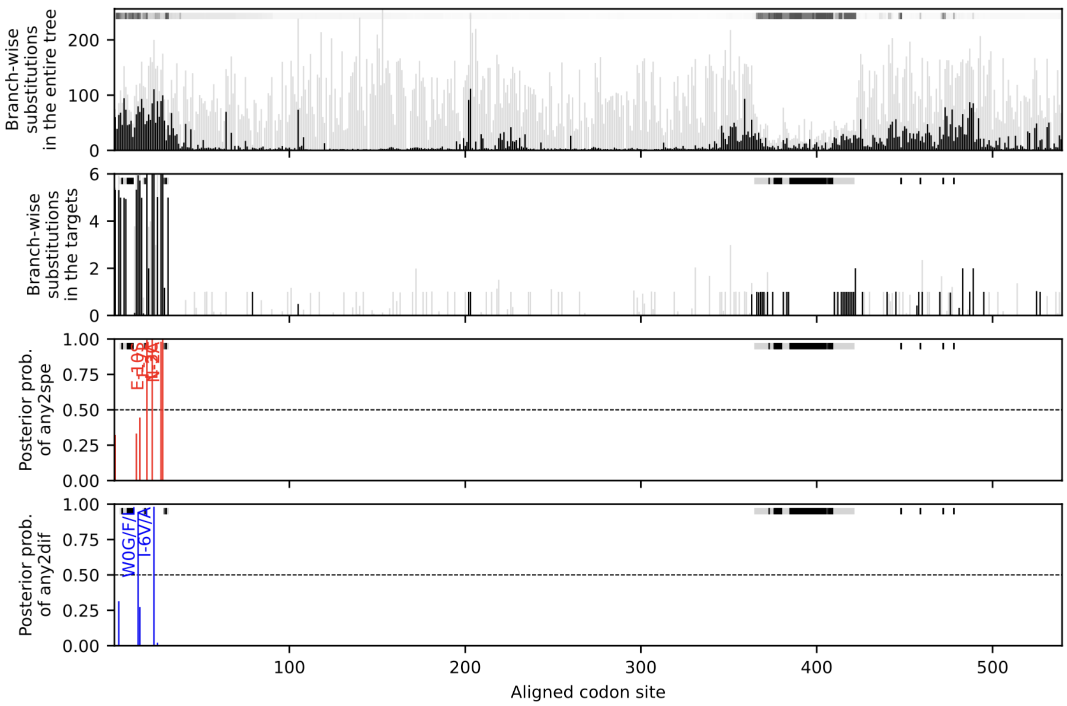
Branch-wise substitutions in the entire tree
Branch-wise substitutions in the targets
Posterior prob. of any2spe
Posterior prob. of any2dif
W0G/F/
I-6V/A
Aligned codon site
0
100
200
300
400
500
0.00
0.25
0.50
0.75
1.00

**B**

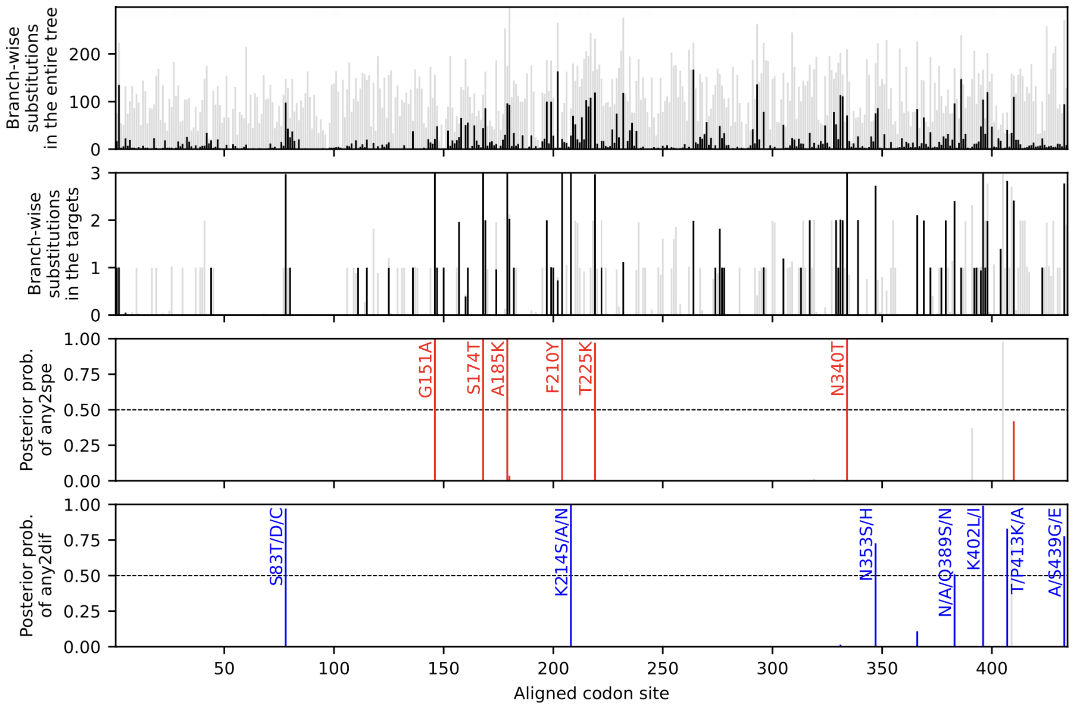
Branch-wise substitutions in the entire tree
Branch-wise substitutions in the targets
Posterior prob. of any2spe
Posterior prob. of any2dif
G151A
S174T
A185K
F210Y
T225K
N340T
S83T/D/C
K214S/A/N
N353S/H
N/A/Q389S/N
K402L/I
T/P413K/A
A/S439G/E
Aligned codon site
50
100
150
200
250
300
350
400
0.00
0.25
0.50
0.75
1.00

**Fig S2: Check for spurious convergence:** There can be certain instances of spurious convergence events even when stringent convergence metrics are used. This can often occur due to misalignment of sequences, which might be due to splice variants represented inconsistently among different species. Fukushima and Pollock also encountered this in their original publication when analysing human and mouse sequences (Supplementary text 12 in (1). This suggests that spurious convergence events are not restricted to any specific dataset but is an artefact that has to be corrected. (A) The characteristic of these spurious convergence events is their unnatural localisation on the protein structures that can be detected in the output of the CSUBST site function. A workaround would be to select convergence events that are not located a proximity to one another. We perform this ad-hoc filtering by selecting orthologs where the individual substitutions have a high OCN value and are well separated on the protein structure. (B) This helps us identify reliable patterns of convergence as shown. In the above plots black and grey vertical bars represent non-synonymous and synonymous substitutions repsectively. The black horizontal bars in panel A represent gaps in mapping the alignment to the protein structure. The posterior probability of *any2spe* represents the site-wise posterior probabilities of a substitution from a differnt acestral amino acid to a specific extant amino acid, i.e. convergent substitutions, while *any2diff* represents a substitution from any ancestral amino acid to a different extact amino acid. These demarkations were used in Fig 2 of the main text.

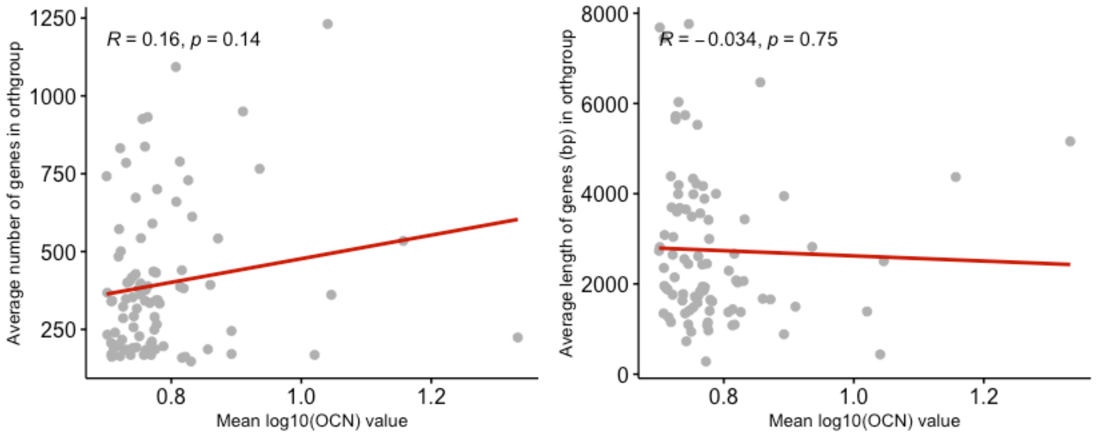


**Fig S3: Observed rate of convergence (OCN) versus orthogroups characterisitcs.** We checked for the presense of any potential bias in between OCN values and the average length and number of genes in each orthogroup. Using a Pearson's correlation test we found no significant relationship between gene number or gene length and the OCN metric, suggesting that the OCN value strictly depends on sequence variation, and that such bias is not a concern in our data.

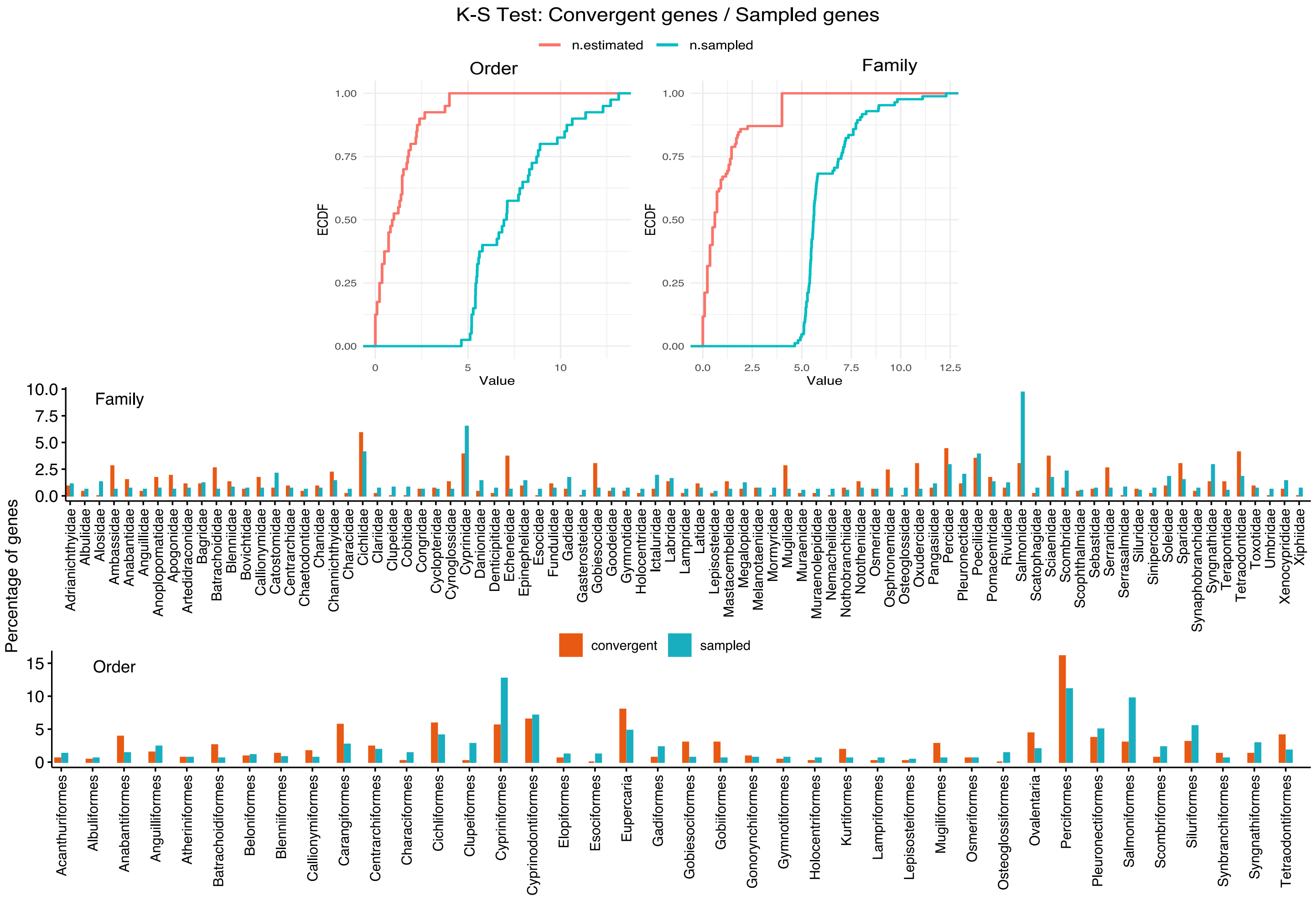


**Fig S4: Comparing the distribution of convergent genes versus sampled genes from each Order.** We use the Kolmogorov-Smirnov test (KS test) to compare distributions of total number of genes sampled versus number of convergent genes estimated (top panle). The rationale is to check whether the relative distributions of the convergent genes are different from that of the number of genes that were sampled. This will tell us whether the deviation between the number of gene sampled in each order/family and the number of genes found convergent is meaningful. The KS test is done with bootstrapping which is suitable for frequency distributions of discrete variables (number of genes in each Order). See https://rdrr.io/cran/kldtools/man/ksboot.html. Observe that the shape of the empirical cumulative probability distributions (ECDF) are different for the convergent genes (red) and total sampled genes (blue). The differences in frequencies is further visualised in the ar plots.

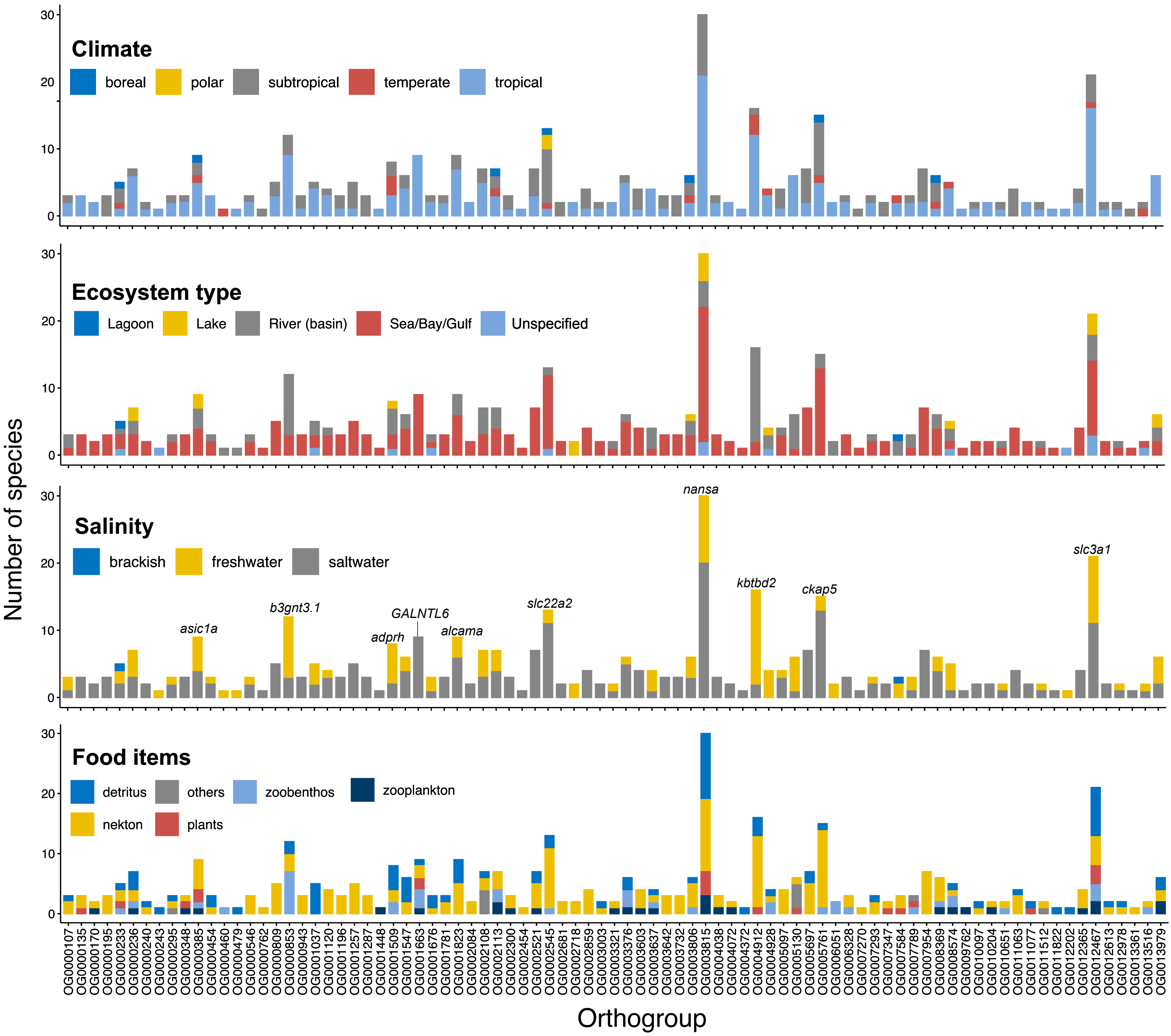


**Fig S5: Ecological characteristics of species with convergent genes.** The bar plots show ecological characteristics of species that harbour convergent substitutions. The data were obtained from the FishBase database. The proportions are based on species with available data. The gene names or orthogroups with a high number of species are labelled. We performed tests of independence to infer whether there is any relationship between convergence in one orthogroup and the ecological characters. After correcting for multiple testing, only 4 orthogroups show a significant relationship. However, these relationships are with the undefined food variable 'other'. As a result, we cannot make any biological relevant inference from this relationship.

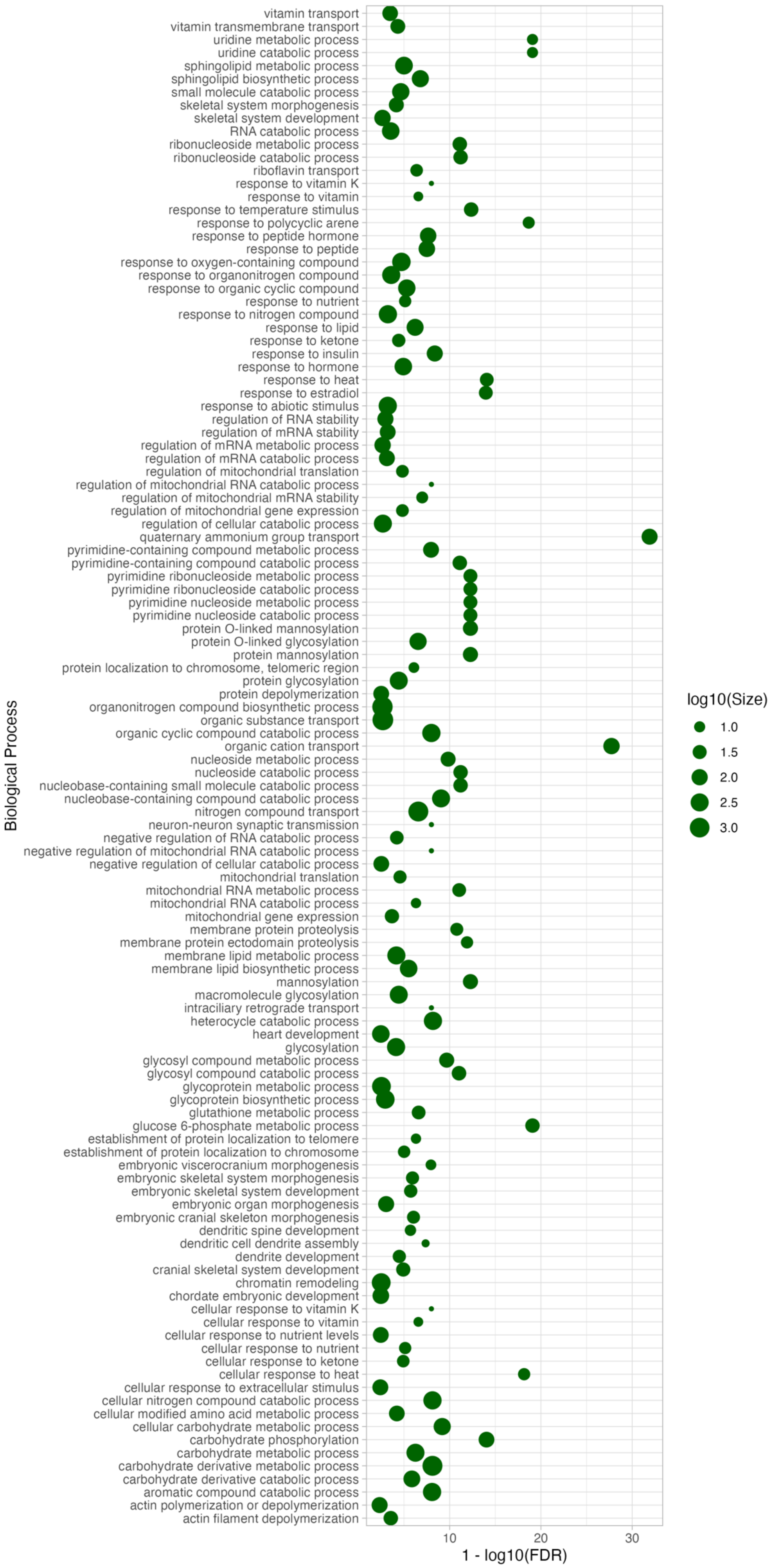
Biological Process
vitamin transport
vitamin transmembrane transport
uridine metabolic process
uridine catabolic process
sphingolipid metabolic process
sphingolipid biosynthetic process
small molecule catabolic process
skeletal system morphogenesis
skeletal system development
RNA catabolic process
ribonucleoside metabolic process
ribonucleoside catabolic process
riboflavin transport
response to vitamin K
response to vitamin
response to temperature stimulus
response to polycyclic arene
response to peptide hormone
response to peptide
response to oxygen-containing compound
response to organonitrogen compound
response to organic cyclic compound
response to nutrient
response to nitrogen compound
response to lipid
response to ketone
response to insulin
response to hormone
response to heat
response to estradiol
response to abiotic stimulus
regulation of RNA stability
regulation of mRNA stability
regulation of mRNA metabolic process
regulation of mRNA catabolic process
regulation of mitochondrial translation
regulation of mitochondrial RNA catabolic process
regulation of mitochondrial mRNA stability
regulation of mitochondrial gene expression
regulation of cellular catabolic process
quaternary ammonium group transport
pyrimidine-containing compound metabolic process
pyrimidine-containing compound catabolic process
pyrimidine ribonucleoside metabolic process
pyrimidine ribonucleoside catabolic process
pyrimidine nucleoside metabolic process
pyrimidine nucleoside catabolic process
protein O-linked mannosylation
protein O-linked glycosylation
protein mannosylation
protein localization to chromosome, telomeric region
protein glycosylation
protein depolymerization
organonitrogen compound biosynthetic process
organic substance transport
organic cyclic compound catabolic process
organic cation transport
nucleoside metabolic process
nucleoside catabolic process
nucleobase-containing small molecule catabolic process
nucleobase-containing compound catabolic process
nitrogen compound transport
neuron-neuron synaptic transmission
negative regulation of RNA catabolic process
negative regulation of mitochondrial RNA catabolic process
negative regulation of cellular catabolic process
mitochondrial translation
mitochondrial RNA metabolic process
mitochondrial RNA catabolic process
mitochondrial gene expression
membrane protein proteolysis
membrane protein ectodomain proteolysis
membrane lipid metabolic process
membrane lipid biosynthetic process
mannosylation
macromolecule glycosylation
intraciliary retrograde transport
heterocycle catabolic process
heart development
glycosylation
glycosyl compound metabolic process
glycosyl compound catabolic process
glycoprotein metabolic process
glycoprotein biosynthetic process
glutathione metabolic process
glucose 6-phosphate metabolic process
establishment of protein localization to telomere
establishment of protein localization to chromosome
embryonic viscerocranium morphogenesis
embryonic skeletal system morphogenesis
embryonic skeletal system development
embryonic organ morphogenesis
embryonic cranial skeleton morphogenesis
dendritic spine development
dendritic cell dendrite assembly
dendrite development
cranial skeletal system development
chromatin remodeling
chordate embryonic development
cellular response to vitamin K
cellular response to vitamin
cellular response to nutrient levels
cellular response to nutrient
cellular response to ketone
cellular response to heat
cellular response to extracellular stimulus
cellular nitrogen compound catabolic process
cellular modified amino acid metabolic process
cellular carbohydrate metabolic process
carbohydrate phosphorylation
carbohydrate metabolic process
carbohydrate derivative metabolic process
carbohydrate derivative catabolic process
aromatic compound catabolic process
actin polymerization or depolymerization
actin filament depolymerization
10
20
30
1 - log10(FDR)
log10(Size)
1.0
1.5
2.0
2.5
3.0

**Fig S6: GO term enrichment of the convergent substitutions.** GO term enrichment of the convergent genes showed that they were inolved in processes related to biomolecule metabolism, response to hormones or stimuli such as heat, and processes of embryonic development and tissue morphogenesis. This underscores the potential multifunctional nature of these convergent genes.

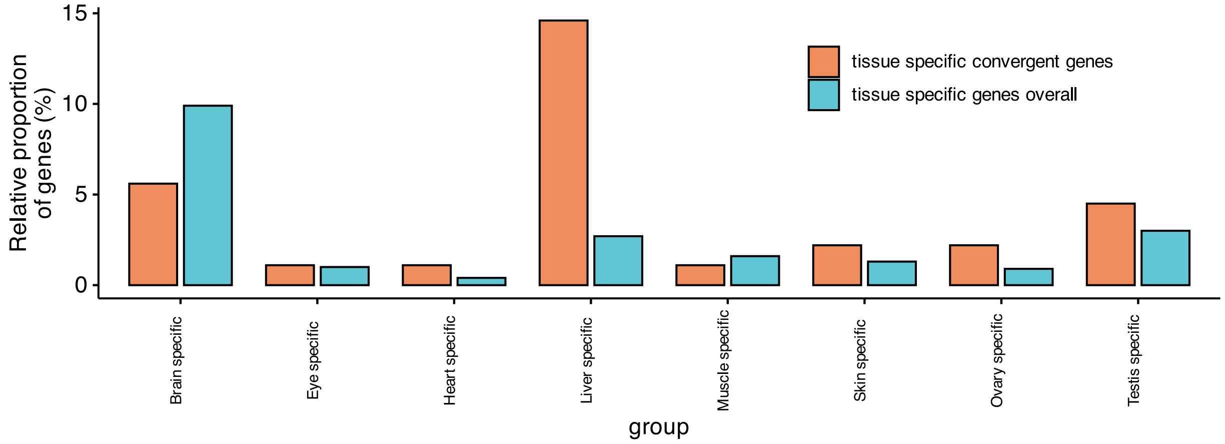


**Fig S7: Only one-third of the convergent genes were tissue specific:** We collected gene expression data for the brain, eye, heart, liver, muscle tissue, ovary, and testis across eleven species (*Astyanax mexicanus, Astatotilapia calliptera, Danio rerio, Esox lucius, Gasterosteus aculeatus, Gadus morhua, Neolamprologus brichardi, Nothobranchius furzeri, Oryzias latipes, Salmo salar,* and *Scophthalmus maximus*). We classified a gene as tissue-specific if it had a tau > 0.8 and its expression in the target tissue was greater than the sum of its expression in other tissues. Since we are comparing tissue specificity across different species, we identified genes that are tissue-specific in the same tissue across all the species sampled. These criteria ensured we captured robust signals for tissue specificity. Around one-third of these genes had signals of convergent evolution with the largest being in the liver. We observed that the proportion of convergent genes that are tissue specific is no different from the total proportion of tissue specific genes in our dataset. However, using Fisher's exact test we found a significant difference in the convergent/non-convergent ratio between tissue-specific and non-tissue-specific genes only for the liver and not for other tissues; in other words, our convergent gene set had a higher proportion of liver-specific genes than expected by chance.

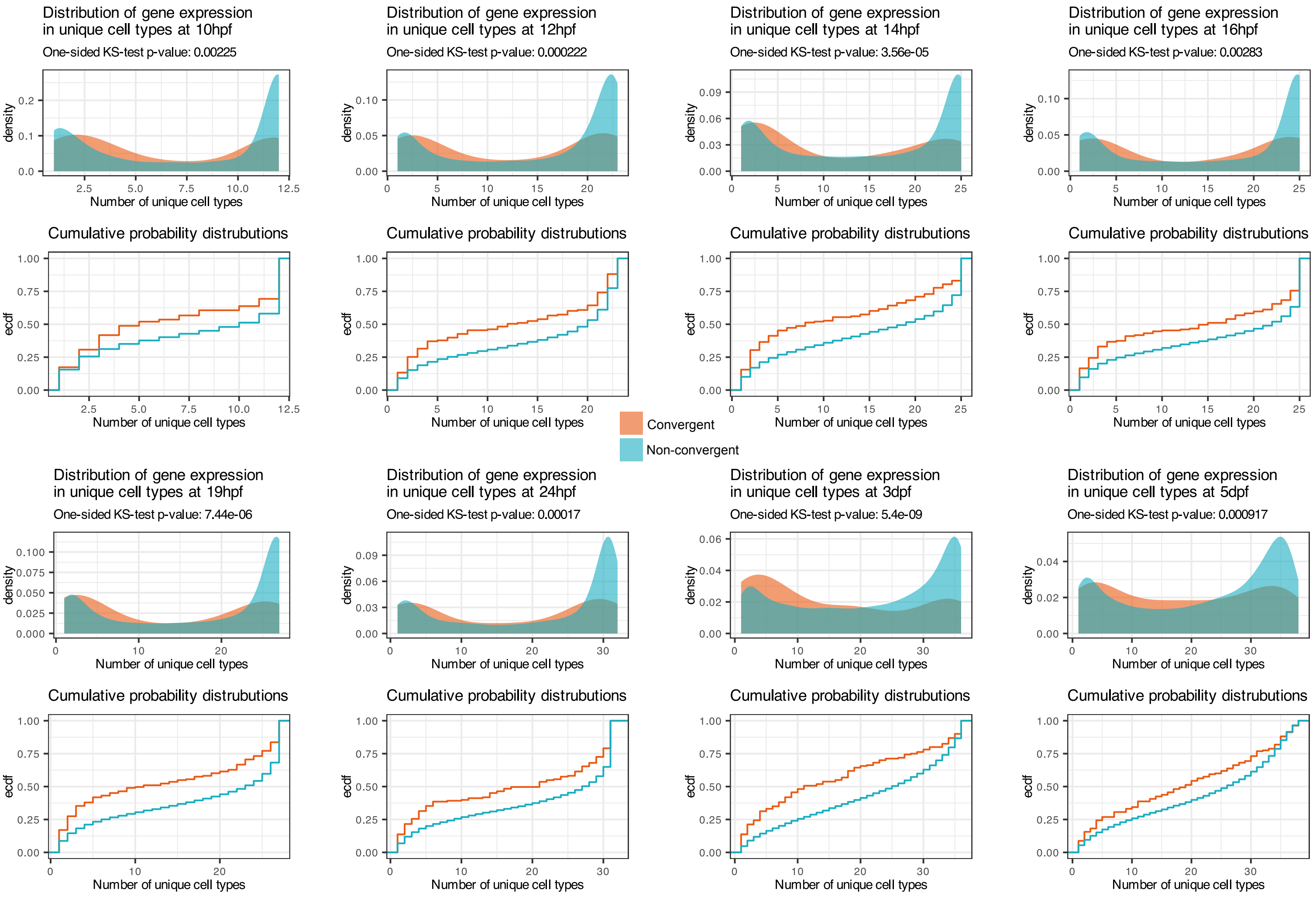


**Fig S8: Distribution of convergent genes at the single-cell level.** At this level we also observed that convergent genes were expressed in an intermediate number of cell types rather than at the extremes of the distribution. The deviations in the cumulative probability distributions are higher at the centre than at the edges of the distribution.

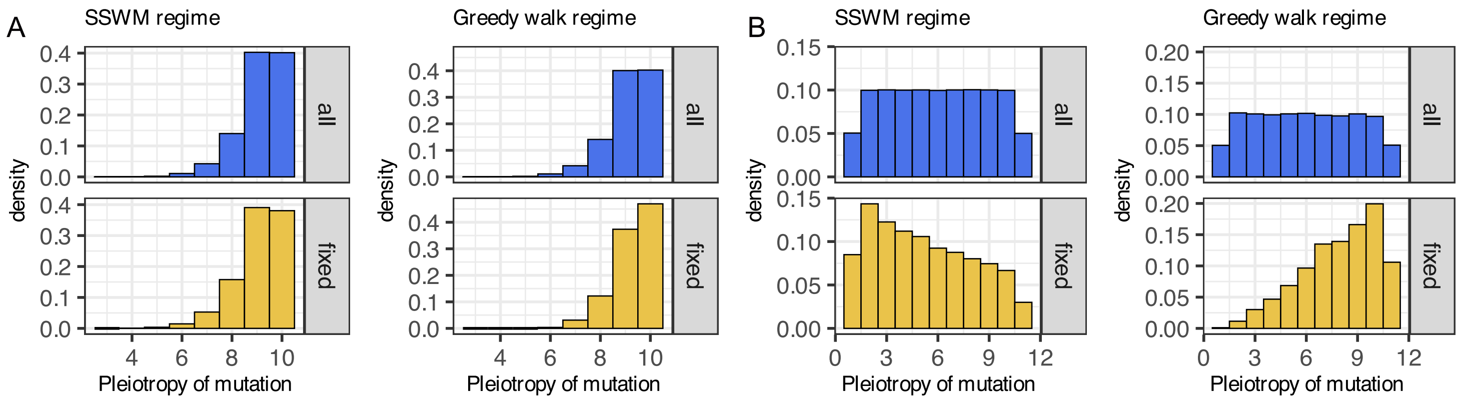


**Fig S9: Artificial pleiotropy distributions used for evolutionary analysis:** To determine the specificity of our model, we also modelled pleiotropy using two artificial distributions: (A) reverse L shape where majority genes have high pleiotropy, and (B) a uniform distribution where all genes have the same pleiotropy. For the reverse L distribution, both regimes had approximately the same distribution of available and fixed mutations. In the uniform distribution, the SSWM regime fixes nearly all available mutations with a slight preference for those with lower pleiotropy. This was expected because, with only a few mutations available, there is no competition between high- and low-pleiotropy mutations. However, since mutations can have either positive or negative fitness effects, lower-pleiotropy mutations, which generally have smaller effects, are more likely to fix, i.e. more mutations with lower pleioytropy mutations tend to be prefered here because they can have lower negative fitness effects. In contrast, the greedy adaptive walk regime shows a strong preference for highly pleiotropic mutations, which was also expected. In this regime, where many mutations are available, those with higher pleiotropy and positive fitness effects tend to fix first. As a result, adaptation in this scenario predominantly occurs through pleiotropic mutations. Fixation patterns using these artificial distributions demonstrate the specificity of the preference of intermediate pleiotropy which is only observed using a real-word empirical distirbution.

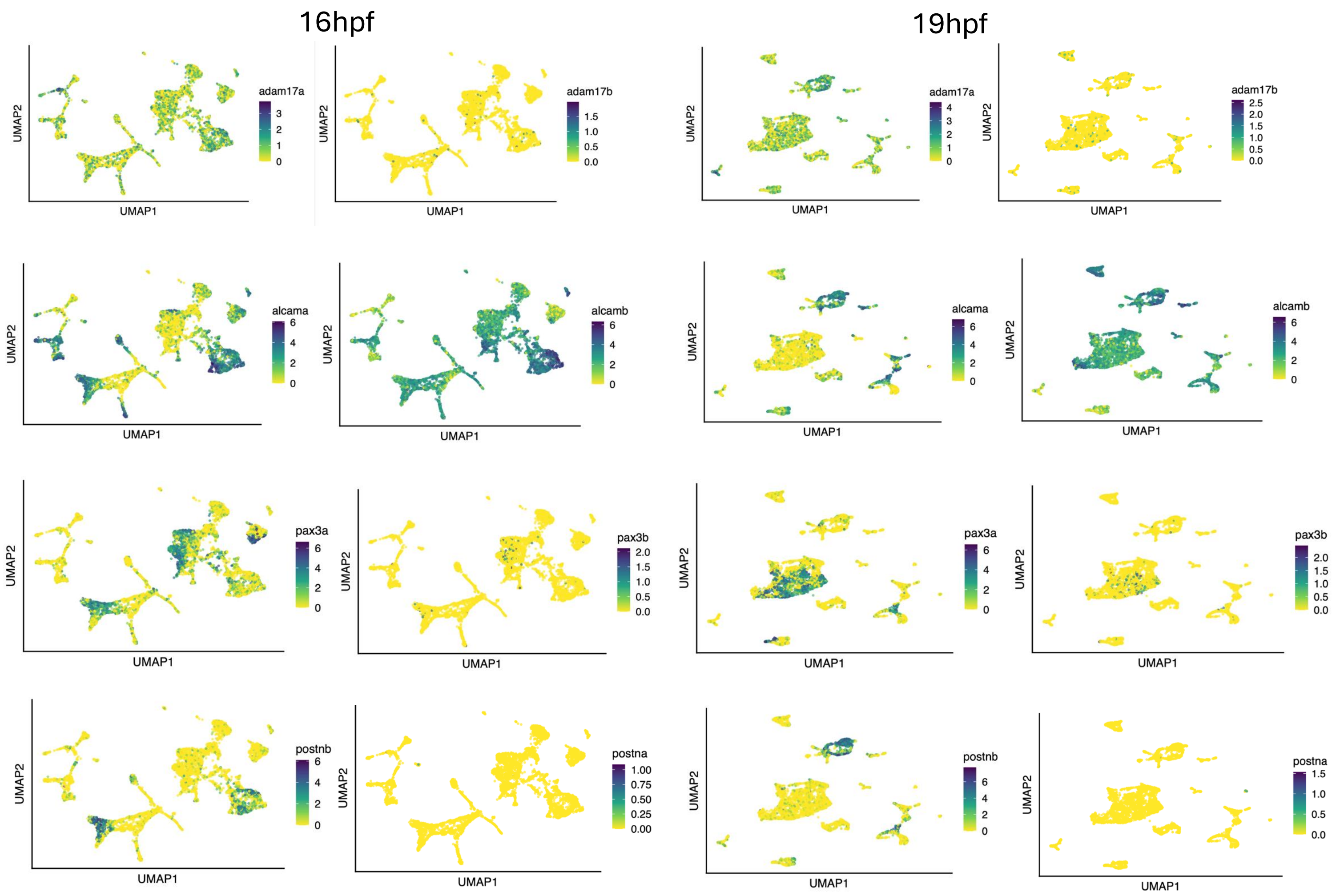


**Fig S10: Expression divergence between paralogs**. Single-cell RNA-seq data of 16 hours post fertilization and 19 hours post fertilization embryos showing difference in gene expression between paralogs. Similar trend was observed for all other time-points.

**Supplementary text 1: Evolution by changes at the protein level**

We observed that convergent substitutions led to changes in several physicochemical properties of amino acids. While experimental validation is necessary—especially regarding protein-protein interactions—the well-documented impact of physicochemical alterations on protein function suggests that these substitutions could have functional consequences[1]. A shift in polarity, for example, can alter hydrophilic or hydrophobic interactions, affecting protein solubility, ligand binding, or membrane interactions. Similarly, mutations that influence secondary structure formation, such as alpha helices, can impact protein stability and function. Changes in steric constraints may disrupt residue positioning, affecting protein dynamics, flexibility, and interactions with other molecules. Several examples exist in the literature of how alterations to the above physicochemical properties, even through a single amino acid change, can lead to global changes in protein activity. One example is the artificially induced L304H mutation in the *VDR* gene in mice (*mVDR*)[2]. This mutation primarily reduces hydrophobicity, increases steric hindrance, and increases polarity, preventing the vitamin D receptor from responding to its natural ligand, $1,25(OH)_2D_3$, while allowing activation by synthetic ligands[2]. Another well-documented case where multiple single substitutions alter physicochemical properties are in the disease-associated mutations at position E22 in amyloid-β peptide (Aβ42), which are linked to Alzheimer's disease[3]. For example, the E22G mutation removes electrostatic repulsion, accelerating aggregation and stabilizing toxic oligomers instead of mature fibrils. The E22K replaces glutamate with lysine, introducing a positive charge. This results in moderate aggregation with lower toxicity. In the E22Q, the change to glutamine, removes the negative charge but maintains polarity. This stabilizes aggregation by promoting hydrogen bonding over ionic interactions, leading to protease-resistant toxic oligomers.

Most of our observed substitutions occur on protein surfaces, where they may influence protein interactions and cellular localisation. Unlike disease-associated mutations, these substitutions could enhance protein-protein interactions, improving specificity and efficiency in biological processes. Over time, they may further optimise interactions, fine-tuning protein function for specialised cellular contexts. Additionally, amino acid changes can reshape protein conformation and tertiary structure, allowing proteins to acquire new functions without disrupting existing catalytic sites[4]. This would enable multifunctionality, affecting different processes across cell types without causing interference. These findings emphasise the importance of considering cellular context alongside genetic changes when studying trait evolution.


1. Kidera, A., Konishi, Y., Oka, M., Ooi, T. & Scheraga, H. A. Statistical analysis of the physical properties of the 20 naturally occurring amino acids. *J. Protein Chem.* **4**, 23–55 (1985).
2. Huet, T. *et al.* A vitamin D receptor selectively activated by Gemini analogs reveals ligand dependent and independent effects. *Cell Rep.* **10**, 516–526 (2015).
3. Kassler, K., Horn, A. H. C. & Sticht, H. Effect of pathogenic mutations on the structure and dynamics of Alzheimer's A beta 42-amyloid oligomers. *J. Mol. Model.* **16**, 1011–1020 (2010).
4. Kozome, D., Sljoka, A. & Laurino, P. Remote loop evolution reveals a complex biological function for chitinase enzymes beyond the active site. *Nat. Commun.* **15**, 3227 (2024).
